# Label-free digital holotomography reveals ibuprofen-induced morphological changes to red blood cells

**DOI:** 10.1101/2022.12.13.519447

**Authors:** Talia Bergaglio, Shayon Bhattacharya, Damien Thompson, Peter Niraj Nirmalraj

**Author notes:** **Author Contributions:** T.B. and P.N.N. planned the study. T.B. prepared the blood samples and conducted the holotomography and AFM measurements, image processing and data analysis. P.N.N. conducted the AFM measurements on ibuprofen particles. S.B. and D.T. conducted the molecular dynamics study and analyzed the simulation results. T.B., S.B., D.T. and P.N.N. wrote the manuscript. All authors discussed the results and commented on the manuscript.

## Abstract

Understanding the dose-dependent effect of over-the-counter drugs on red blood cells (RBCs) is crucial for hematology and digital pathology. Yet, it is challenging to continuously record the real-time, drug-induced nanoscopic shape changes of RBCs in a label-free manner. Here, we demonstrate digital holotomography (DHTM) enabled real-time, label-free concentration-dependent and time-dependent monitoring of ibuprofen on RBCs from a healthy donor. The RBCs are segmented based on 3D and 4D refractive index tomograms and their morphological and chemical parameters are retrieved with their shapes classified using machine learning. We directly observed the formation and motion of spicules on the RBC membranes when aqueous solutions of ibuprofen were drop cast on wet blood, creating rough-membraned echinocyte forms. At low concentrations of 0.25-0.50 mM, the ibuprofen-induced morphological change was transient but at high concentrations (1.5-3 mM) the spiculated RBC remained over a period of up to 1.5 hours. Molecular simulations confirmed that aggregates of ibuprofen molecules at high concentrations significantly disrupted the RBC membrane structural integrity and lipid order, but produced negligible effect at low ibuprofen concentrations. Control experiments on the effect of urea, hydrogen peroxide and aqueous solutions on RBCs showed zero spicule formation. Our work elucidates the dose-dependent chemical effects on RBCs using label-free microscopes that can be deployed for the rapid detection of overdosage of over-the-counter and prescribed drugs.

**Significance:** The interaction between drugs and blood cells is an important field of study in order to understand the risk for drug-induced haematological adverse effects. Using digital holo-tomographic microscopy (DHTM), we can resolve the real-time effect of medications on the morphological and chemical properties of red blood cells with high spatial and temporal resolution and in a label-free manner. We show that our approach can be used as a haematology platform for the diagnosis of blood disorders and for monitoring the dose-dependent effect of prescribed and over-the-counter medications in a cost-effective manner, with significant implications for its applicability in resource-limited settings and in the field of personalized medicine.

## Introduction

The rheological properties of RBCs including deformability and aggregability help regulate blood flow through the circulatory system (1, 2). Impaired RBC deformability can result in increased blood viscosity, impaired perfusion, occlusions in small blood vessels and could lead to ischemia (3, 4). Other factors can trigger changes to the membrane mechanical properties of RBCs, concentration changes of hemoglobin inside the cell and modifications of the RBC surface area or volume (5–8). These factors range from the primary genetic mutations in the different forms of hereditary hemolytic anemia to secondary processes arising from mechanical or chemical alterations in the surrounding environment. The overall RBC shape change is conventionally used to describe RBC deformability. Failure to maintain optimal red cell deformability results in a lower RBC life span and contributes to the development of hemolytic anemia (6). One pathology resulting from hemolytic anemia is sickle cell disease (SCD), a group of inherited hematological disorders affecting hemoglobin (9). In sickle cell anemia (SCA), a significant population of RBCs are shaped as sickles, thus becoming less deformable, with a lower life span and with an increased risk of blood clot formation, infections and pain (9, 10). Interactions with drugs (for example, vinblastine, colchicine and chlorpromazine) and signaling molecules can also negatively influence red cell rheological properties, by decreasing RBC deformability and increasing RBC aggregation (2, 3, 11).

Hemolytic anemia can thus be induced by a wide range of medications including cephalosporin-based antibiotics and oxaliplatin anti-cancer drugs (12–14). Despite the low incidence rate of this side effect, drug-induced immune hemolytic anemia (DIIHA) is a serious condition most observed from the use of OTC medications due to the higher risk of their misuse (12, 15). Among these medications are nonsteroidal anti-inflammatory drugs (NSAIDs) widely used for their anti-inflammatory, antipyretic and analgesic properties (16). Ibuprofen is one such NSAID used for the treatment of rheumatoid arthritis and for the relief of pain, inflammation and fever (15). In addition to increased risk of gastrointestinal injury, ibuprofen can influence hemostasis even at recommended doses, causing thrombocytopenia, reduced platelet aggregation resulting in increased clotting time and loss of hemoglobin, potentially leading to DIIHA (15–17). NSAID toxic side effects may result from their interaction with cellular membranes, which primarily act as a protective barrier and regulate materials transfer into and out of the cell, including drug delivery, based on precise molecular-level organization, fluidity and permeability (17, 18). The RBC membrane can be considered an ideal model for the investigation of drug–cell interaction due to the presence of a single phospholipid bilayer membrane and the absence of internal organelles inside RBCs (11, 18, 19).

Hence, monitoring RBC deformability constitutes a crucial diagnostic tool, with shape change patterns allowing patient stratification by disease stage, and can be used in a pre-clinical setting to gauge the effect of pharmacological interventions on blood-related disorders such as SCD, thalassemia, diabetes and COVID-19 (6, 20–23). Despite the well-reported interaction between ibuprofen and the lipid bilayer membrane (19), to the best of our knowledge only one study has shown the potential effects of ibuprofen on RBC deformability (17). In this study, Manrique *et al*. (17) used scanning electron microscopy (SEM) to obtain snapshots of RBC morphological changes after incubation at different concentrations of ibuprofen and provide evidence for spicule formation on the cell membrane and echinocytosis with increasing ibuprofen dosages. Importantly, although RBC shape changes were observed with ibuprofen concentrations as low as 10 μM, the reversibility of these changes could not be determined due to the lack of dynamic cell behavior information. Hence, the effect of ibuprofen on RBC morphology and the dose-dependent interactions with the RBC lipid bilayer membrane remains to be clarified with high spatial clarity in real-time.

Label-free digital holotomographic microscopy (DHTM) enables 3D morphometric imaging of live cells with nanoscale resolution at room temperature (24). Unlike fluorescent microscopy, DHTM does not rely on fluorescent labelling and uses a low-power laser beam that avoids phototoxic effects (24). Several studies have demonstrated the potential of digital holography in the field of hematology (25–29). In particular, Kim *et al*. have previously demonstrated the use of common-path diffraction optical tomography (cDOT) for the visualization of RBCs and the quantification of RBC morphometric parameters (28). Moreover, holotomography was employed to study the mechanobiology of RBCs upon exposure to Melittin (30). Unlike previous studies, here we directly register the dosage dependent effect of ibuprofen on RBCs in real-time using DHTM.

We demonstrate here a DHTM based approach for label-free detection and quantification of ibuprofen-induced RBC shape changes with high spatial (~200 nm) and temporal resolution (~2 s). As a control, we recorded the DHTM RI maps of RBCs from healthy individuals and from those with sickle cell anemia (SCA) and sickle cell trait (SCT) condition. The 3D and 4D RI tomograms were analyzed using a machine learning (ML) based classifier to identify difference in shapes between RBCs in healthy and pathological individuals (SCA and SCT). Next, we extended the imaging and analytics protocols to investigate the concentration and time dependent effect of ibuprofen on RBCs from healthy individuals. Monitoring the real-time changes in RBC morphology upon ibuprofen introduction from 0 to 20 minutes, we observed the formation of spicules on the RBC membrane, defined as echinocytosis. The nanoscopic details of the spicule morphology was further analyzed using atomic force microscopy (AFM) and the real-time motion of the spicules on RBC membrane was captured using DHTM. Spicule formation was observed to be reversible at lower ibuprofen concentrations (0.25 mM and 0.5 mM), but the normal RBC discocyte morphology did not recover with higher ibuprofen concentrations (1.5 mM and 3 mM), over a period of up to 1.5 hours. To understand the interaction and effect of ibuprofen molecules on RBC membrane morphology at experimentally inaccessible timescales of molecule– molecule interaction (0-100 ns), we conducted atomic-scale molecular dynamics (MD) computer simulations. Models of membrane-bound single, very small (n=80), small (n=100), and large (n=1903) aggregates of ibuprofen molecules confirmed the extensive deformation of RBC lipid bilayer only at high concentrations with large aggregates of ibuprofen. Further control experimental measurements of drug-free RBCs in water and other chemicals such as urea and hydrogen peroxide (H_2_O_2_) confirmed that spicule formation only occurs with ibuprofen. These findings suggest that high-throughput microscopy and ML-driven automated image analysis methods provide a valuable platform for the early diagnosis of blood disorders and for monitoring the efficiency of prescribed and OTC drugs in a simple, field-deployable and cost-effective manner (20, 31).

## Results

### Quantification and classification of RBCs using label-free DHTM

In order to deduce the chemical effects on RBCs, we first compared samples from a healthy donor, a donor diagnosed with sickle cell trait (SCT) and a donor diagnosed with sickle cell anemia (SCA). This benchmarking of the DHTM tool enabled live cell and artefact-free imaging. The principle and experimental scheme of DHTM is explained in Fig. 1*A*. DHTM allows for the fast acquisition of refractive index (RI) tomograms rendered in 3D that provide quantitative information regarding RBC morphology. Details on the preparation of RBC samples for DHTM imaging are provided in *Materials and Methods* (Fig. 1*B*) and the demographic information on donors is provided in *SI Appendix*, Table S1. Fig. 2*A* shows a 3D RI tomogram of healthy RBCs diluted in phosphate-buffered saline (PBS) solution, with a field of view of 90×90×30 μm. Here, the distinctive biconcave disciform shape of an RBC can be observed, with the inner part of the cell having a lower RI value compared to the outer area due to the concavity of the disk shape (27, 28). Fig. 2*B* shows the corresponding segmented RI tomogram with the background signal removed and the voxels extracted for the single RBCs. The RBCs could be classified based on their morphology using a ML-based algorithm. As shown in Fig. 2*C*, all cells shown in the image were classified as normocytes. The same approach was applied to RBCs extracted from donors with SCT (Fig. 2*D-F*) and SCA (Fig. 2*G-I*). A greater variability in RBC morphology was evident in both SCT and SCA samples, including the presence of echinocytes, acanthocytes, spherocytes and sickle RBCs (Fig. 2*F* and *I*). Importantly, the observed variability in cell morphology could at least in part be attributed to the transport and long storage time (~15 days) between blood collection and analysis of the SCT and SCA samples. Long storage periods can negatively influence RBC rheological properties by altering RBC morphology from discocytes to echinocytes, creating a potential confounding effect in the assessment of RBC health, particularly in individuals with a blood-related pathology characterized by RBC morphological alterations, such as SCA (32, 33). For example, we observed from control experiments that blood from a healthy donor, diluted in PBS and stored at 4°C over a period of 11 days, resulted in the morphological transition of RBCs from mostly normocytes on day 0 to an increasing population of echinocytes over normocytes up to 70% on day 11 (*SI Appendix*, Fig. S1).

**Fig. 1.**
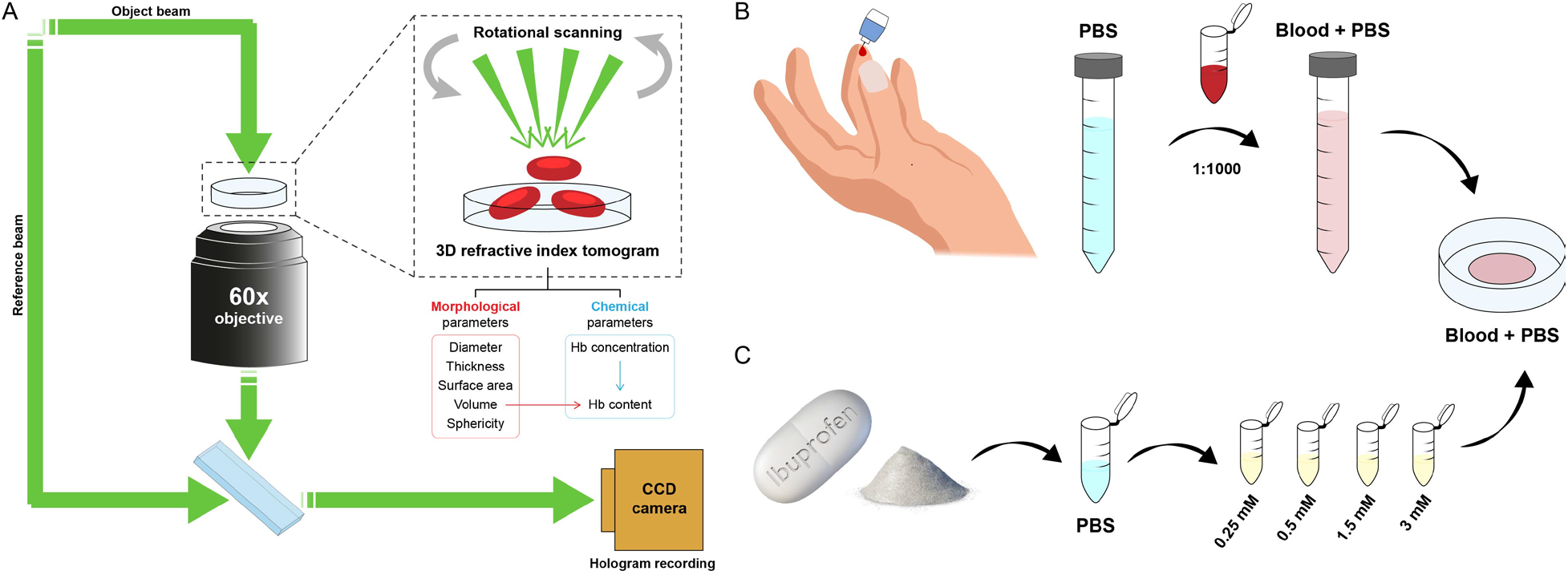
Principle of DHTM and sample preparation procedure for blood and ibuprofen solutions. (*A*) The holo-tomographic setup includes a low power laser beam (λ = 520 nm) that splits into the reference and the object beam before rejoining below the objective, where the interference is recorded. A 3D RI map is obtained by recording holograms with a rotational arm at 360° around the sample, at a 45° angle. Morphological and chemical parameters can be quantified for individual RBCs from the 3D RI tomogram. (*B*) 10 μL of whole blood is obtained from a finger prick and diluted in PBS buffer at a final concentration of 1:1000. 250 μL of blood solution is added to a petri dish for imaging. (*C*) Ibuprofen powder is obtained by crushing an ibuprofen tablet and is dissolved in PBS buffer to obtain four final concentrations (0.25 mM, 0.5 mM, 1.5 mM, 3 mM). 50 μL of each ibuprofen solution is added to the blood solution in the petri dish during the live cell imaging experiments.

**Fig. 2.**
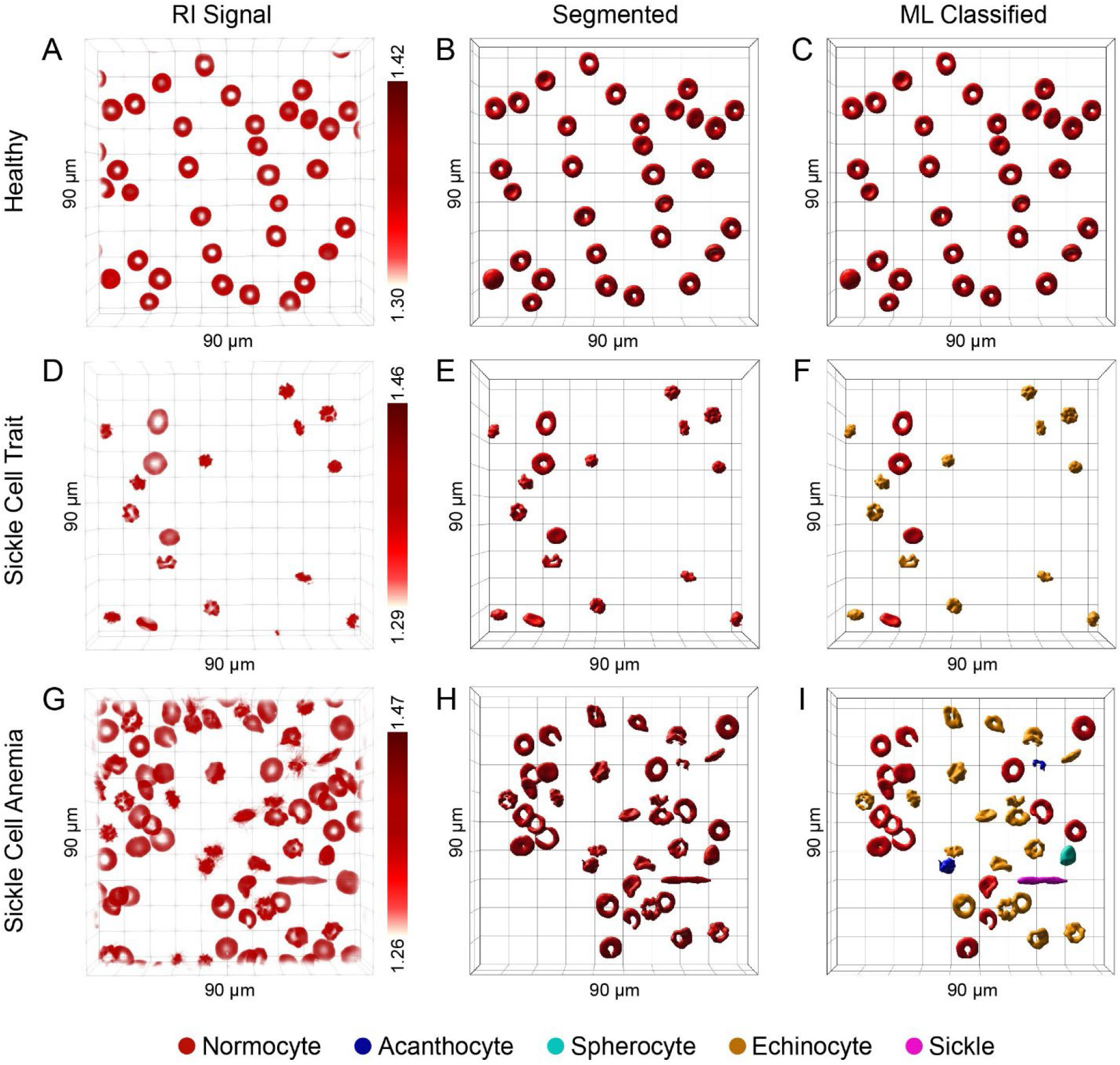
3D holotomographic imaging of RBCs and classification using machine learning (ML). (*A*) 3D refractive index (RI) tomogram of RBCs obtained from a healthy donor. The corresponding segmented RI tomogram and ML-classified RBC types are shown in (*B*) and (*C*). (*D*) 3D RI tomogram of RBCs from a donor with sickle cell trait (SCT). The corresponding segmented RI tomogram and ML-classified RBC types are shown in (*E*) and (*F*). (*G*) 3D RI tomogram of RBCs from a donor diagnosed with sickle cell anemia (SCA). The corresponding segmented RI tomogram and ML-classified RBC types are shown in (*H*) and (*I*). Field of view 90×90×30 μm.

From the segmented RI tomograms, the quantitative information of the RBC morphological and chemical parameters, including diameter, surface area, volume, thickness, sphericity, hemoglobin (Hb) concentration and Hb content, was extracted at a single cell level (Fig. 3). In order to assess the accuracy of the quantification of the RBC morphological measurements based on DHTM RI tomograms, we used microparticles of silicon dioxide with nominal sizes of 2 and 5 μm (*SI Appendix*, Fig. S2). The Imaris-based segmentation and quantification method yielded similar results in terms of bead diameter, surface area and volume compared to the nominal values reported by the manufacturer. A total of 351 healthy, 459 SCT and 230 SCA RBCs were analyzed and classified into RBC types (*SI Appendix*, Table S2). The measured values for normocytes were comparable in both healthy, SCT and SCA samples and were consistent with the literature, with a reported mean diameter of 8 μm (Fig. 3*A*), mean surface area of 130 μm^2^ (Fig. 3*B*) and mean volume of 90 fL (Fig. 3*C*) (34). Fig. 3*A* reveals variations in mean diameter between different RBC types. Stomatocytes (healthy = 6.88 μm, SCT = 6.18 μm, SCA = 7.95 μm), echinocytes (healthy = 6.69 μm, SCT = 6.56 μm, SCA = 7.63 μm), acanthocytes (SCT = 5.60 μm, SCA = 8.31 μm) and spherocytes (SCT = 5.03 μm, SCA = 6.16 μm) had a lower diameter compared to normocytes (healthy = 7.77 μm, SCT = 7.81 μm, SCA = 8.45 μm). Conversely, sickle RBCs found in SCT (8.52 μm) and SCA (12 μm) samples had a higher diameter due to their elongated shape compared to normocytes. The values for the mean surface area (Fig. 3*B*) and mean volume (Fig. 3*C*) showed corresponding lower values for echinocytes (healthy: 100 μm^2^, 77.7 fL; SCT: 91.64 μm^2^, 59.28 fL; SCA: 115.79 μm^2^, 75.20 fL), acanthocytes (SCT: 72.35 μm^2^, 43.94 fL; SCA: 110.24 μm^2^, 74.10 fL) and spherocytes (SCT: 64.46 μm^2^, 45.27 fL; SCA: 79.78 μm^2^, 52.84 fL) compared to normocytes (healthy: 127.51 μm^2^, 96.29 fL; SCT: 119.75 μm^2^, 79.53 fL; SCA: 129.58 μm^2^, 86.88 fL). Spherocytes found in SCT and SCA samples also showed a slightly higher thickness (SCT = 2.26 μm, SCA = 1.77 μm) compared to normocytes (SCT = 2.03 μm, SCA = 1.56 μm) due to the transition from a biconcave disciform shape to a spheroid morphology (Fig. 3*D*). Consequently, the same pattern was found for the sphericity morphological parameter (Fig. 3*E*), with values closer to 1, indicating a perfect sphere, (SCT = 0.94, SCA = 0.85) compared to normocytes (SCT = 0.75, SCA = 0.73). A similar pattern was observed in terms of thickness (Fig. 3*D*) and sphericity (Fig. 3*E*) for both echinocytes (healthy: 2.21 μm, 0.88; SCT: 1.77 μm, 0.81; SCA: 1.67 μm, 0.75) and acanthocytes (SCT: 1.83 μm, 0.83; SCA: 1.47 μm, 0.78) due to the tendency of these RBC types to be more spherical in shape compared to normal RBCs. The results from DHTM and our analysis methodology indicate that the RI can be used as a metric assessing Hb concentration and Hb content, as the cytoplasm of RBCs contains mainly Hb solution (see *Materials and Methods* for details on the calculation of Hb concentration and Hb content from RI values) (29). Based on DHTM measurements, we observed a slightly higher Hb concentration and a corresponding lower Hb content (*SI Appendix*, Fig. S3) for all RBC types compared to normocytes, which we attribute to the changes in RBC shape, specifically a decrease in RBC volume, and thus the possible rearrangement of Hb within a single RBC (Fig. 3*F*). The retrieved mean Hb concentration (healthy = 35.2 ± 0.5 g/dL, SCT = 35.5 ± 1.2 g/dL, SCA = 34.4 ± 0.7 g/dL) and mean Hb content (healthy = 33.9 ± 4.0 pg, SCT = 28.1 ± 5.3 pg SCA = 29.9 ± 4.4 pg) for normocytes are in agreement with the reference values for the mean corpuscular Hb concentration (MCHC) and the mean corpuscular Hb (MCH) reported in a complete blood count (CBC) of healthy individuals (MCHC = 32-36 g/dL, MCH = 28-32 pg) (35).

**Fig. 3.**
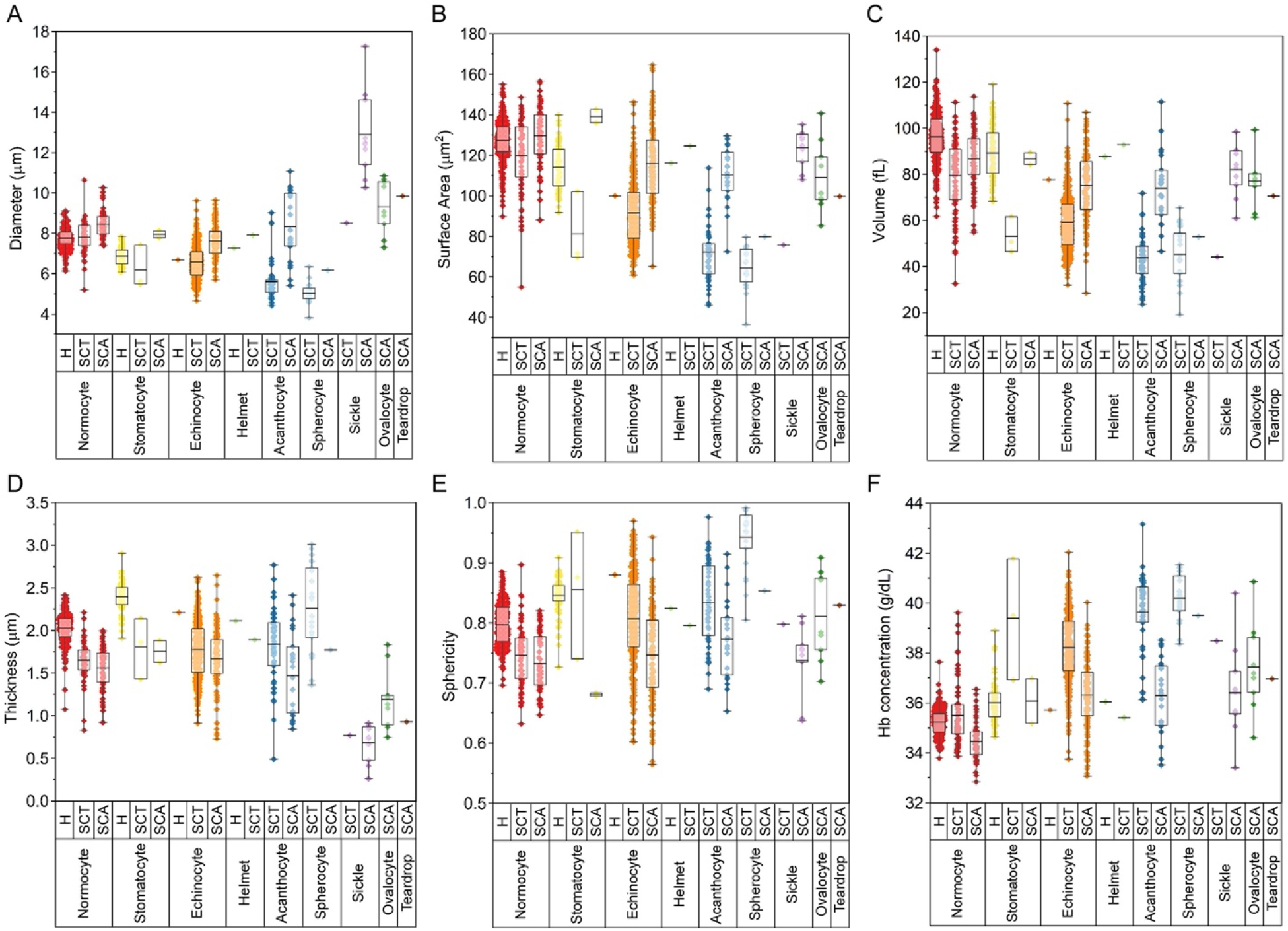
Quantification of size and shape variations in healthy, SCT and SCA RBC populations based on 3D tomograms. Single cell level comparison of (*A*) diameter, (*B*) surface area, (*C*) volume, (*D*) thickness, (*E*) sphericity and (*F*) Hb concentration between ML-based classified RBC types in healthy, SCT and SCA samples. Bars indicate mean values plus minimum and maximum values of all counted cells in each group.

With the imaging and analysis framework described in our study, we were able to optimize the DHTM based imaging technique to accurately resolve and quantify single RBCs in a label-free manner in both healthy and disease states. In addition, using a ML-based classification approach, we were able to distinguish between different RBC types and identify morphological and chemical parameters that could be used to describe changes in RBC shape, as benchmarked against previous studies on RBC morphology using other label-free imaging methods (7, 20, 22, 26, 28).

### Dose-dependent effect of ibuprofen on RBCs

The dose-dependent effect of ibuprofen on healthy RBCs was evaluated in real-time using DHTM and analyzed using the protocols described above for the comparative analysis of RBCs from healthy, sickle cell trait and sickle cell anemia donors. Fig. 4 shows the segmented and ML-classified RI tomograms of RBCs during incubation with different concentrations (0.25 mM-3 mM) of ibuprofen, over a period of 20 minutes. For all ibuprofen concentrations, the formation of spicules on the RBC membrane and a clear transition from normocytes to echinocytes was observed upon introduction of ibuprofen in the RBC environment (> 34s). At low ibuprofen concentrations (0.25 mM and 0.5 mM) (Fig. 4*A-B*), the loss of the normal biconcave disciform morphology was determined to be transient, with most RBCs transitioning from normocytes to echinocytes and back to normocytes within 20 minutes (*SI Appendix*, Movie S1-S2). However, at high ibuprofen concentrations (1.5 mM and 3 mM) (Fig. 4*C-D*), the echinocytosis deformation did not result in the recovery of the normocyte RBC morphology (*SI Appendix*, Movie S3-S4). In order to further investigate the dynamics of spicule formation, movement and ultimately dissolution across the RBC membrane, we imaged and quantified single cell dynamics for low and high ibuprofen concentrations (0.25 mM and 1.5 mM) (Fig. 5). As shown in fig. 5*A*, the segmented 3D single RBC begins transitioning into an echinocyte upon exposure to 0.25 mM of ibuprofen (t = 44 s), with spicules forming on the RBC membrane, and continues to dynamically change before returning to a normal biconcave disciform shape at the 20 minutes time point. During this time, spicules can be observed forming (t = 44 s and 1:14 min), merging (t = 7:04 min), splitting (t = 7:36 min) and finally dissolving (t = 20 min) (*SI Appendix*, Movie S5). The corresponding variations in RI, as shown in the insets in fig. 5*A*, reveal a rearrangement of hemoglobin inside the cell during the morphological transition, with areas containing protrusions having a higher RI value (1.39) compared to flatter regions (1.33). The time-dependent reversible morphological changes of the single RBC were quantified and are shown in fig. 5*B-E*. Upon the introduction of ibuprofen, the cell diameter (Fig. 5*B*) decreased from 7.39 μm to as low as 6.57 μm and was followed by a gradual increase back to 7.87 μm at 20 minutes, associated with the transition from a normocyte to an echinocyte shape and later returning to a discocyte morphology. Likewise, the surface-area to volume (S/V) ratio suffered an initial drop from 1.36 to 1.20, driven by a decrease in surface area unmatched by a decrease in cell volume (*SI Appendix*, Fig. S4*A-B*), that later recovered up to 1.28. Upon the transition to a more spherical echinocyte-shaped RBC, the cell sphericity (Fig. 5*D*) also increased up to 0.90 and gradually returned within the range of a normocyte. Hb concentration (Fig. 5*E*) was also subjected to a transient increase upon the introduction of ibuprofen, as reflected by the RI map inset in Fig. 5*A*. The higher values represent the time of spicule formation and movement across the RBC membrane, and the values gradually returning to a slightly lower Hb concentration compared to the starting point. Hb content initially decreased upon the introduction of ibuprofen and later returned to slightly higher values by the 20 minutes time point (*SI Appendix*, Fig. S4*C*).

**Fig. 4.**
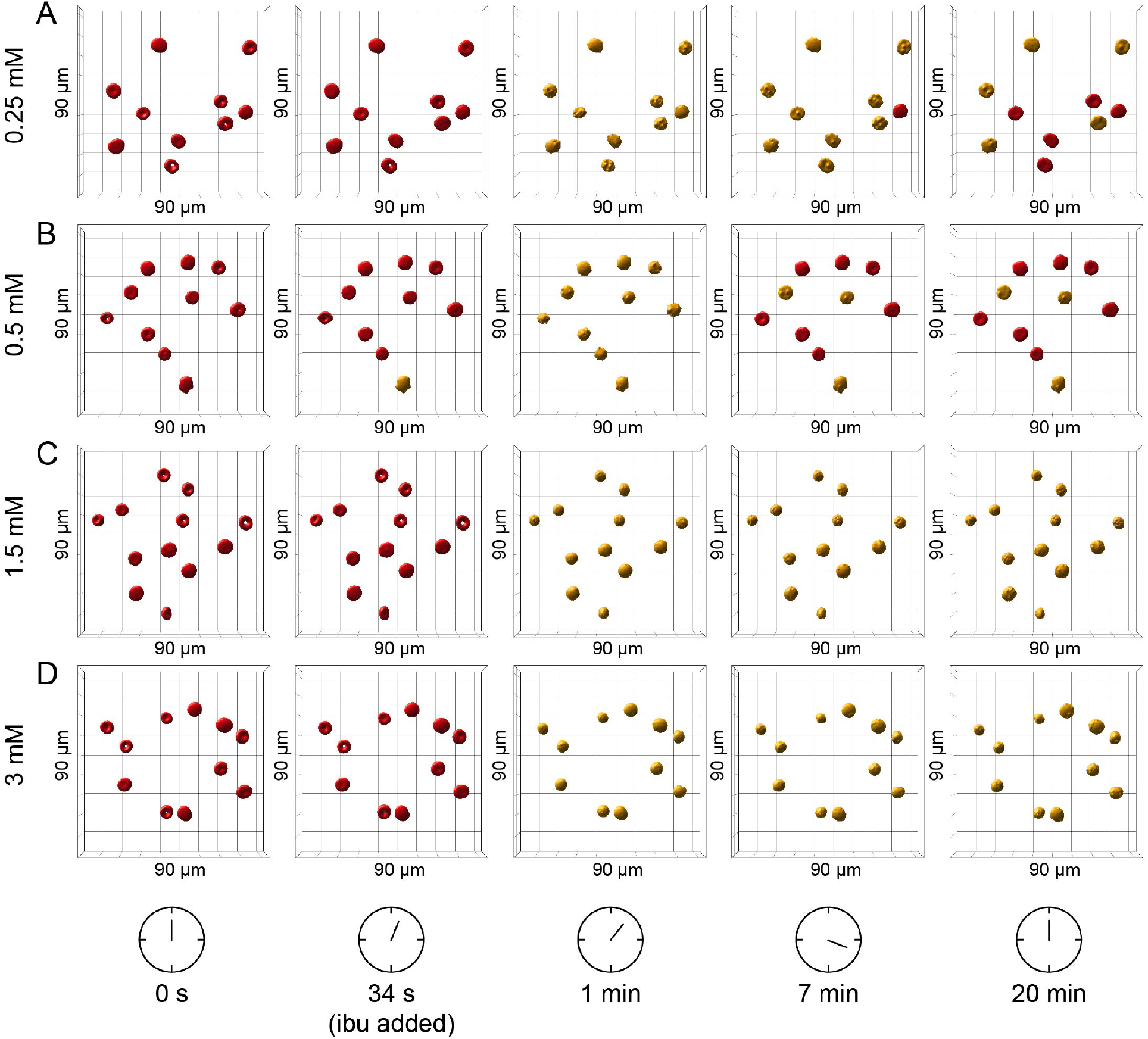
3D rendering and classification of RBCs treated with ibuprofen at varying concentrations during a 20-minute time-lapse using 3D digital holo-tomographic microscopy. (*A*) 0.25 mM, (*B*) 0.5 mM, (*C*) 1.5 mM, and (*D*) 3 mM. Red and yellow color coding indicates normocytes and echinocytes, respectively. Field of view 90×90×30 μm. ibu = ibuprofen.

**Fig. 5.**
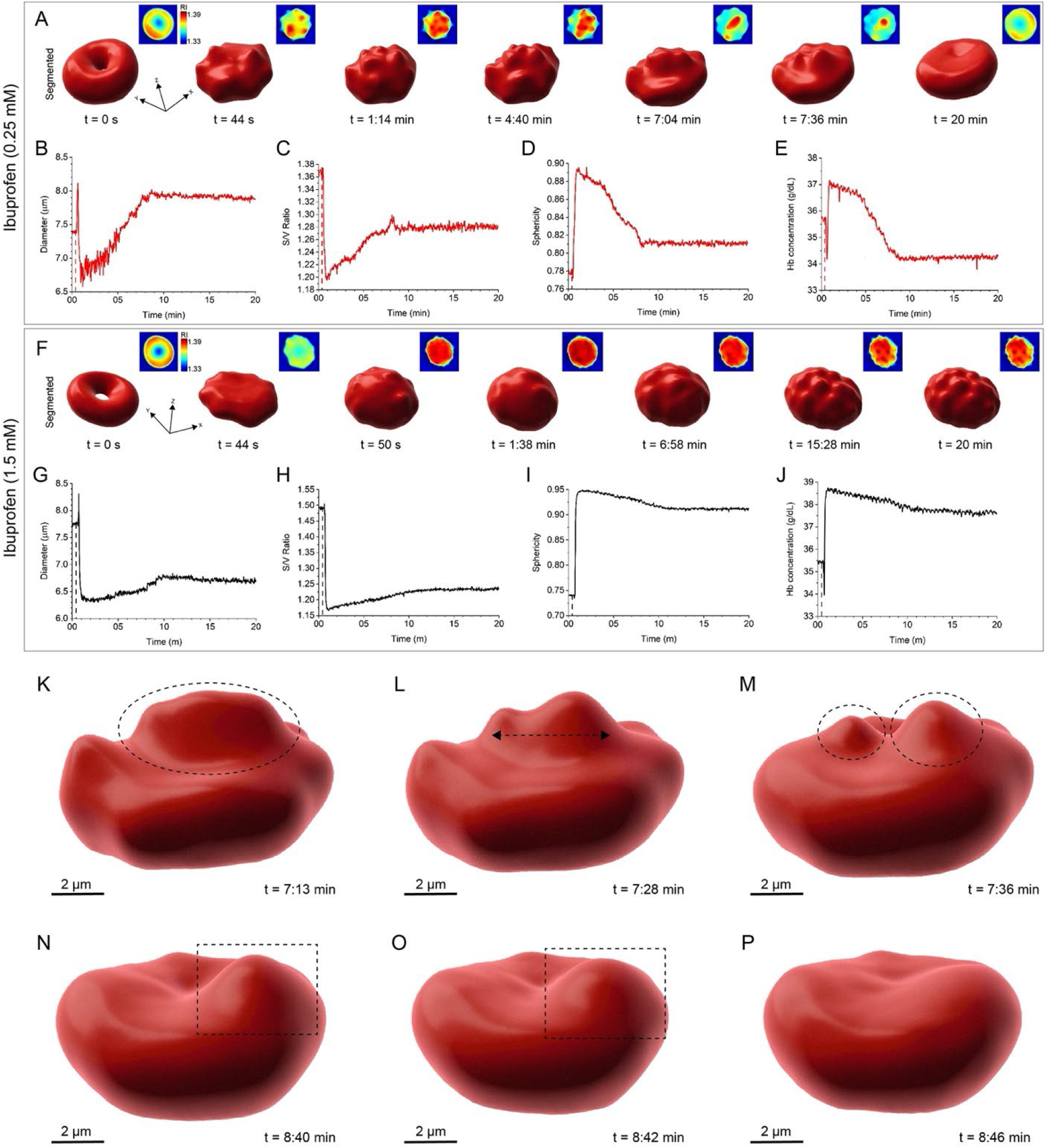
RBC morphological changes upon exposure to low and high concentrations of ibuprofen during a 20-minute time-lapse. (*A*) 3D renderings of a single RBC treated with 0.25 mM ibuprofen showing the transient morphological alteration from a normocyte to an echinocyte (scale bar: *x* = 7.57 μm, *y* = 7.36 μm, *z* = 3.28 μm). Insets in (*A*) show the corresponding 2D RI maps at each time point. (*B*)-(E) Quantification of time-dependent morphological parameters: diameter, S/V ratio, sphericity and Hb concentration. (*F*) 3D renderings of a single RBC treated with 1.5 mM ibuprofen showing sphero-echinocytosis (scale bar: *x* = 7.74 μm, *y* = 6.92 μm, *z* = 2.82 μm). Insets in (*F*) show the corresponding 2D RI maps at each time point. (*G)-(J*) Quantification of time-dependent morphological parameters: diameter, S/V ratio, sphericity and Hb concentration. (*K-M*) 3D rendering of a single RBC treated with 0.25 mM ibuprofen showing spiucle splitting. (*N-P*) 3D rendering of a single RBC treated with 0.25 mM ibuprofen showing spicule dissolution.

The segmented 3D individual RBCs treated with 1.5 mM ibuprofen is shown in Fig. 5*F* and the quantified RBC parameters are shown in fig. 5*G-J*. At high ibuprofen concentration, spicules were observed forming and slightly moving across the RBC membrane but never dissolved by the 20 minutes time point (*SI Appendix*, Movie S6). Upon exposure to ibuprofen, the RBC transitioned to a sphero-echinocyte with a lower diameter, from 7.74 μm to 6.69 μm (Fig. 5*G*), a reduced S/V ratio ranging from 1.49 to 1.24 (Fig. 5*H*) with decrease in surface area unmatched by decrease in cell volume (*SI Appendix*, Fig. S4*D-E*), and a significant increase in cell sphericity (Fig. 5*I*) up to 0.95 and later of 0.91 at the 20 minutes time point, reaching values very close to the sphericity of a perfect sphere. The increase in Hb concentration (Fig. 5*J*) and Hb content (*SI Appendix*, Fig. S4*F*) after exposure to high ibuprofen concentrations was associated with the morphological transition to a sphero-echinocyte, with the most significant protrusions showing the highest RI values (Fig. 5*F* inset). Spicule movement on the RBC membrane was observed with high resolution and at a single cell level for all ibuprofen-treated RBCs. An example of a spicule that splits into two daughter spicules within a ~20 s time period is shown in fig. 5*K-M*. Similarly, the dynamic dissolution of a spicule was observed in RBCs treated with low ibuprofen concentration and is portrayed in fig. 5*N-P*.

For the nanoscopic characterization of spicules on the RBC membrane, we used AFM in tapping mode to analyze RBCs present in air-dried blood. The blood smears were prepared after incubation of healthy blood with different ibuprofen concentrations (0.25 mM, 0.5 mM, 1.5 mM, 3 mM) for up to 1.5 hours (see *Materials and Methods* for details on sample preparation). A progressive increase in the number of echinocytes over normocytes was observed with increasing ibuprofen concentrations, with the majority of the RBCs incubated with 3 mM ibuprofen maintaining the echinocyte morphology after 1.5 hours (*SI Appendix*, Fig. S5). In view of the 1-2 hours half-life of ibuprofen (36), we suggest that with a high ibuprofen concentrations (3 mM), echinocytosis is likely to persist even when ibuprofen has been excreted. Fig. 6*A* shows a 3D rendered AFM height image of a single echinocyte RBC, with the white arrows indicating the individual spicules on the RBC membrane. The corresponding height and phase-contrast AFM images are shown in fig. 6*B-C*. Variations in height between the flatter regions of RBC compared to the regions containing protrusions are visible in the AFM topograph shown in fig. 6*A* and were quantified using line sectional analysis as shown in fig. 6*D*. Compared to the height profile extracted from a healthy normocyte shown in green and in the inset in fig. 6*D*, the height profile of the echinocyte (blue line) presents protrusions of variable sizes, ranging from ~100 to 300 nm, and does not show the typical biconcave disciform profile. Analysis of surface roughness, normalized over an RBC area of 1 μm^2^, between ibuprofen-treated RBCs and healthy RBCs revealed a stark difference, with a higher mean surface roughness of 40.5 ± 20.8 nm for ibuprofen-treated RBCs compared to 8.9 ± 6.6 nm for healthy RBCs. Qualitative and quantitative variations in RBC morphology and membrane topography are clearly distinguishable between ibuprofen-treated RBCs and normocytes. In order to register the size of ibuprofen aggregates, ibuprofen solution (concentration: 9.7 mM) was deposited on a gold thin film and the particles were measured using AFM. Fig. 6*F* shows a height AFM image of the ibuprofen particles. The ibuprofen aggregates were measured on an atomically clean gold surface (surface roughness: <0.5 nm) instead of directly on the RBC surface because the surface of RBCs could also contain other protein aggregates, even in healthy donors, which can result in the misleading estimation of ibuprofen particle size distribution (37). Averaging over the result from several line sectional profiles similar to those shown in Fig. 6G, we calculate a mean ibuprofen particle size of 13.5 nm, with confidence interval lower (CIL) bound of 11.5 nm and confidence interval upper (CIU) of 14.9 nm. The confidence interval was calculated at 95% as shown in the non-gaussian statistical distribution plot (Fig. 6H). The quantitative assessment of ibuprofen particle size distribution suggests that ibuprofen drug molecules could be mostly present in the form of aggregates on the surface of RBCs.

**Fig. 6.**
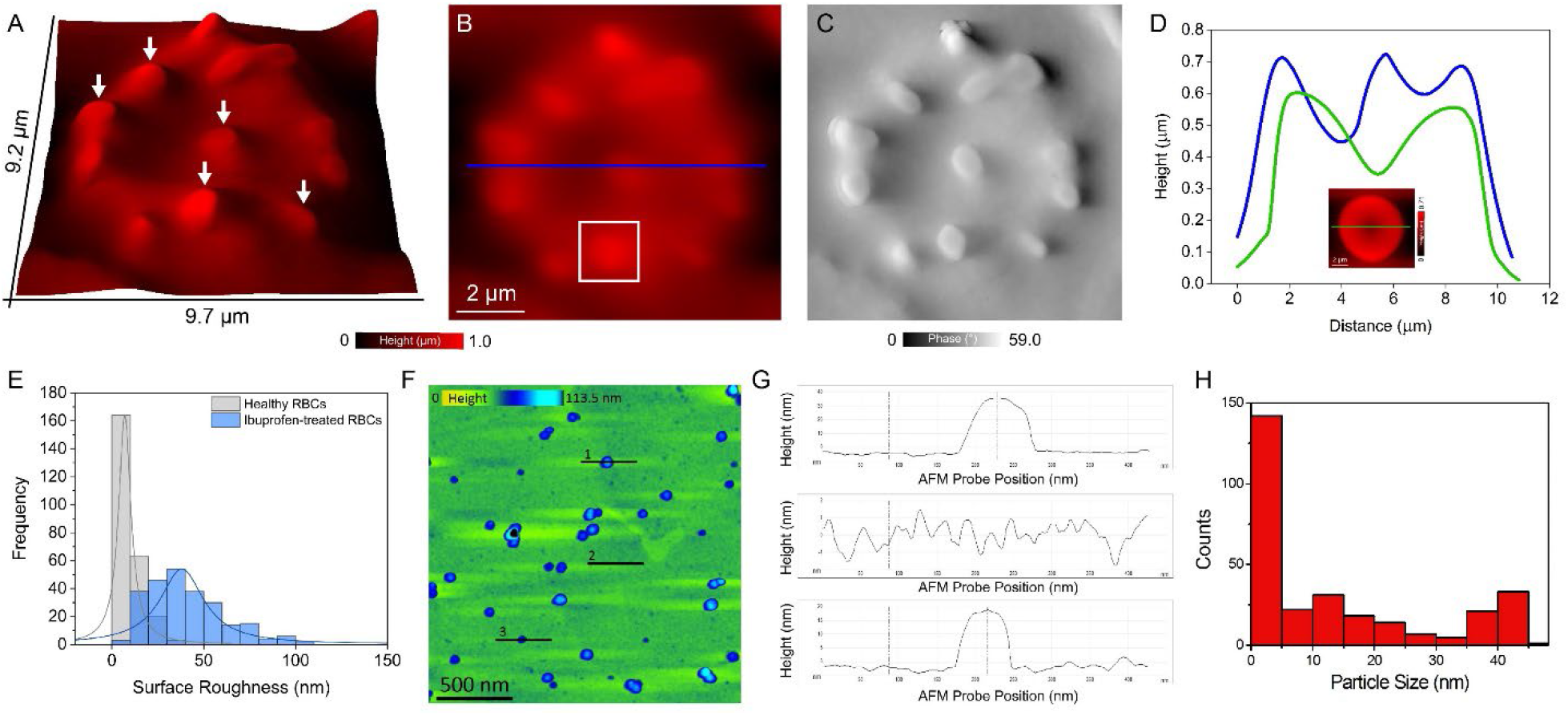
Characterization of spicules on ibuprofen-treated RBCs. (*A*) 3D AFM image of an echinocyte. White arrows indicate spicules on RBC membrane. (*B*) and (*C*) Height and phase-contrast AFM images of an echinocyte. (*D*) Height profile extracted along the blue line indicated in (*B*) across an echinocyte. The green line indicates the corresponding height profile of a normocyte taken from the inset in (*D*). (*E*) Statistical distribution of surface roughness values obtained from AFM based analysis of healthy and ibuprofen-treated RBCs. (F) AFM height image of ibuprofen particles on gold surface. (*G*) Height profiles of ibuprofen particles extracted along the corresponding lines indicated in (*F*). (*G*) Ibuprofen particle size distribution; mean particle size: 13.5 nm, CIL: 11.5 nm and CIU: 14.9 nm.

### Control experiments to study the effect of other chemicals on RBC morphology

In order to verify that spicule formation is a result of ibuprofen treatment, we performed additional control experiments to investigate the effect of drug-free solutions of urea (*SI Appendix*, Fig. S6), H_2_O_2_ (*SI Appendix*, Fig. S7) and double-distilled water (ddH_2_O) (*SI Appendix*, Fig. S8) on RBC morphology. Urea is known to cross the red cell membrane and to weaken the membrane cytoskeleton by perturbing the structure of spectrin (8). RBCs treated with 2 M, 4 M and 6 M urea transitioned into spherocytes, with a lower diameter and an increased thickness, S/V ratio and sphericity (*SI Appendix*, Fig. S6*A-C, E-J*). When 8 M urea was added to RBCs, spherocytosis was followed by vesiculation and lysis, associated with a drop in Hb concentration and Hb content, within ~2 minutes (*SI Appendix*, Fig. S6*D, E-J*). Additionally, the effect of oxidative damage on RBC membrane function was assessed by introducing different concentrations of H_2_O_2_ (2 M, 4 M, 6 M, 8 M) into the RBC environment, which resulted in a transient morphological transformation into stomatocytes. The stomatocytes displayed slightly decreased diameter and Hb concentration, markedly lower sphericity values and a higher S/V ratio, that later mostly recovered to their original normocyte shape at the 15 minutes time point (*SI Appendix*, Fig. S7). H_2_O_2_ has been previously reported to impair RBC deformability by inducing oxidation of hemoglobin, alterations to membrane proteins and lipid peroxidation (38). RBCs treated with ddH_2_O did not undergo any significant morphological change within ~10 minutes, as reflected by a constant diameter, S/V ratio, sphericity and Hb concentration (*SI Appendix*, Fig. S8). Based on these findings, echinocytosis was not observed as a result of urea, H_2_O_2_ and ddH_2_O treatment. In contrast, a morphological transition from normocytes to echinocytes was observed when imaging the RBCs in a petri dish with a glass surface (*SI Appendix*, Fig. S9) and when storing blood diluted in PBS and stored at 4°C over a period of 11 days, in agreement with previous findings, highlighting the importance of studying freshly-collected RBCs for the assessment of RBC morphology (*SI Appendix*, Fig. S1) (32, 33).

Taken together, the DHTM and AFM measurements provide evidence in support of a dose-dependent and time-dependent effect on the ibuprofen induced changes to RBC morphology. Our qualitative and quantitative data confirms that the RBC membrane undergoes distinctive changes when interfacing with ibuprofen drug molecules that can ultimately affect RBC morphology and RBC rheological properties.

### Modelling the effect of low *vs*. high concentrations of ibuprofen on the RBC membrane structure

To understand the experimental dose-dependent effect and interactions of ibuprofen with the RBC membrane lipid bilayer (see Section S1.1, Fig. S11*A*, and Table S4 in *SI Appendix* for details of the membrane model composition), we performed extensive molecular dynamics (MD) computer simulations at different ibuprofen concentrations: one molecule of ibuprofen, which we name “single ibu”; preformed aggregates of 80 ibuprofen molecules, “low ibu conc. I”; 100 ibuprofen molecules, “low ibu conc. II”; densely packed 1903 ibuprofen molecules under constant pressure, “high ibu conc. I”; and 1903 molecules at constant volume, “high ibu conc. II” (see *SI Appendix*, Section S1.2 for more details on the computational models and methods). The structures formed during 0.1 μs of equilibrated, unconstrained dynamics for each system reveal that a single molecule of ibuprofen quickly permeates the RBC lipid outer layer (Fig. S12*F*) and remains bound in the lipid core. This is reflected in the improvement in ibu–lipid interaction energies after 10 ns (Fig. S12*K*) facilitated by favorable hydrophobic van der Waals (vdW) interactions of the ibuprofen propyl tail with the lipid aliphatic chains. The small aggregates of ibuprofen at low ibu conc. I and II make only transient interactions with the membrane bilayer (Fig. S12*G*, *H*, *L, M*) and remain in strongly aggregated clusters driven by ibuprofen–ibuprofen hydrophobic forces (Fig. S13*A*, *B*). Despite the favorable ibuprofen–ibuprofen vdW interactions (Fig. S13*C*, *D*), at the high ibu conc. I and II, the densely packed ibuprofen shows significantly improved interactions with the lipid bilayer (Fig. S12*N*, *O*), leading to disruption of the RBC bilayer as described below.

Computed density profiles of all species (Fig. S13*E*–*I*) show that the thickness of the lipid bilayer is ~7 nm for all but the high ibu conc. II, where the bilayer is compressed to ~6 nm (Fig. S13*I*). The small dip in the water density profile marks the position of the aggregated ibuprofen at low concentrations (Fig. S13*F*, *G*). By contrast, the water density is significantly replaced by densely packed ibuprofen near the outer membrane leaflet at high concentrations (Fig. S13*H*, *I*), also facilitating diffusion of several ibuprofen molecules into the membrane. There is apparent lateral diffusion of lipid molecules across the membrane as evident from the flattening density of the membrane center at high ibu conc. II (Fig. S13*I*), which otherwise shows a dip in membrane density at high ibu conc. I (Fig. S13*H*). This indicates the presence of hydrophobic tails of each leaflet facing each other sampling a dissipated central membrane thickness. To confirm lateral diffusion of lipids in the membrane due to high concentrations of adsorbed ibuprofen, we computed the mean square displacements (MSD) and diffusion coefficients (*D*) of lipid headgroup atoms (P, N, and O) for each system. The MSD plots reveal an increased displacement of lipid headgroups mediated by ibuprofen aggregates, but a significantly larger correlation of MSD with simulation time at high ibu conc. II (Fig. S14*A*). Similarly, the *D* reveals clear distinction between high ibu conc. II and other systems of aggregated ibuprofen on membrane, the former showing a significantly higher diffusivity of the membrane polar headgroups (Fig. S14*B*).

We further mapped the lipid hydrocarbon tail deuterium order parameters (S_CD_, Fig. S14*C*–*G*) showing significant loss of lipid order at high ibu conc. II (see Fig. S14*G*). Finally, we mapped the lipid heavy atoms number densities in the *xy*-plane and averaged over the *z*-axis to obtain a top view of lipid densities in the membrane (Fig. S14*H-L*). An ibuprofen concentration-dependent loss of lipid structuring could be observed where a single ibuprofen does not affect the lipid density (Fig. S14*H*), while at low concentrations, an uneven distribution is revealed (Fig. S14*I*, *J*) with low-density pockets that are most prominent at high ibu conc. II (Fig. S14*L*). Overall, our modelling data predict that at low concentration, ibuprofen does not affect the RBC membrane structure (Fig. 7*A*), but at high concentration, the lipid membrane is deformed (Fig. 7*B*), due to large-area ibuprofen and lipid membrane interaction at high concentration (Fig. 7*C*) driven by hydrophobic vdW forces (see Fig. S12*L*, *O* in SI). The significantly higher lipid diffusion coefficient computed at high ibuprofen concentration reveals that the lipids are in constant motion, while at low concentration, the polar headgroups are more stable (Fig. 7*D*). The disruption of lipid structural integrity at high concentration is supported also by the disordering of acyl carbon atoms (Fig. 7*E*). Finally, lipid number density in the plane of the membrane clearly show the dense and ordered lipid packing at low ibuprofen concentration, as opposed to the non-uniform lipid distribution when highly concentrated densely packed ibuprofen is adborbed on the RBC membrane (Fig. 7*F*). The data suggests that the lipid molecules undergo a substantial RBC membrane morphological deformation when exposed to high doses of ibuprofen but experience little to no change at low ibuprofen doses.

**Fig. 7.**
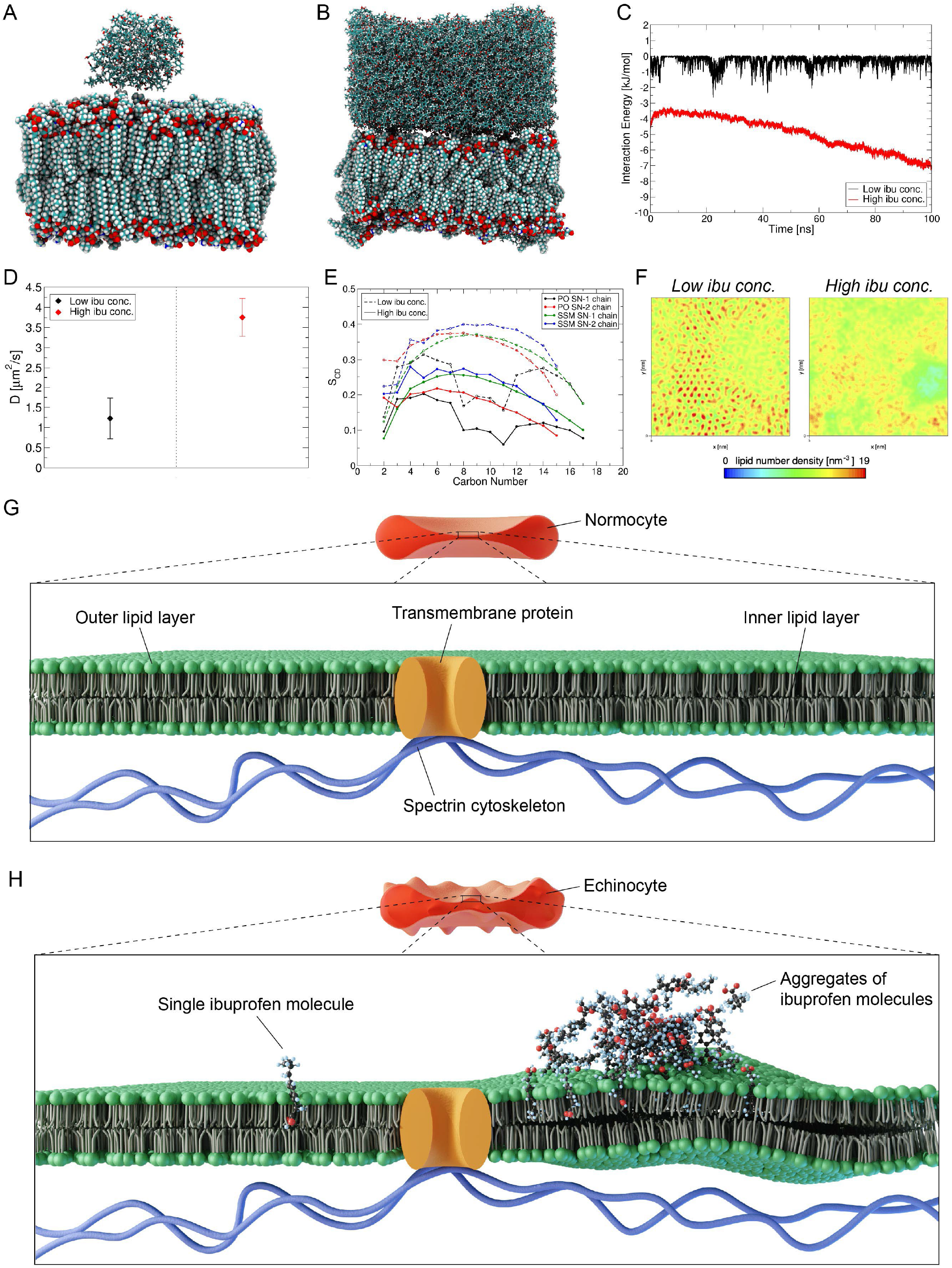
Representative structures of ibuprofen (ibu) aggregates on RBC outer membrane bilayer obtained from Molecular Dynamics (MD) simulations of (*A*) low concentration of ibuprofen adsorbed on membrane, and (*B*) densely packed high concentration of ibuprofen adsorbed on membrane. (*C*) Total interaction energies between ibuprofen and lipid membrane at low and high concentrations. (*D*) Comparison of the diffusion coefficient, D of RBC lipid headgroup atoms (P, N and O) at low (black) and high (red) concentrations of ibuprofen on membrane. (*E*) Deuterium order parameters (S_CD_) of the hydrocarbon tails, SN-1 and SN-2 of Palmitoyl Oleoyl (PO) and Stearoyl (SSM) lipids of the RBC membrane bilayer in presence of low and high concentrations of ibuprofen. (*F*) Maps of average (over the *z*-axis) lipid number density in the plane (*xy*) of the RBC membrane bilayer. (*G*) Schematic representation of a normocytic RBC membrane architecture showing the lipid bilayer, a transmembrane protein and the spectrin cytoskeleton. (*H*) Schematic representation of an echinocyte membrane architecture showing the interaction of one ibuprofen molecule (left) and multiple aggregates of ibuprofen molecules (right) with the lipid bilayer. A single ibuprofen molecule permeates and interacts with the RBC lipid outer layer while bigger ibuprofen molecule aggregates diffuse and deform the lipid bilayer, causing spicule formation.

## Discussion

In this study, we investigated the ibuprofen-induced morphological alterations to RBCs in real-time and in a label-free manner using DHTM. From the 3D RI tomograms, we tracked the formation of spicules on the RBC membrane associated with a clear morphological transition from normocytes to echinocytes upon exposure to ibuprofen drug solutions. The morphological changes in the RBCs were observed to be concentration-dependent and were either transient, at 0.25-0.50 mM ibuprofen concentrations, or never recovered their original shape, at 1.5-3 mM ibuprofen concentrations, monitored over a period of 20 minutes. The RBC morphological parameters were extracted from 3D RI tomograms and quantified as first demonstrated for healthy, SCT and SCA blood samples. The extracted quantitative information on ibuprofen-treated RBCs supported the qualitative evidence. All RBCs exposed to ibuprofen exhibited a decrease in diameter and S/V ratio, which is driven by a lower cell surface area and volume. This change is associated with the transition from normocytes to echinocytes, with simultaneous increase in sphericity and Hb concentration in response to the decrease in RBC volume. Spicules were observed to form, merge, split and dissolve on the RBC membrane, correlating the cell shape alterations with the progression of both echinocytosis and spherocytosis processes. Both the cell parameters and shape of RBCs exposed to low ibuprofen concentrations (equivalent to a 200 mg and 400 mg tablet) gradually recovered after ~8 minutes from the introduction of ibuprofen particles, suggesting a reversible drug-induced effect on the RBC membrane. Previously, echinocytosis has also been observed using cell imaging techniques and attributed to presence of excessive EDTA, prolonged storage of RBCs prior to preparation of blood smears on solid surfaces and pathological causes such as in liver and kidney diseases (39, 40). However, in the present study we attribute the formation of spicules to the interaction of ibuprofen at high concentrations with RBCs. We deduce this result based on label-free imaging of healthy RBCs interacting with ibuprofen, urea (2M-8M), H_2_O_2_ (2M-8M) and ddH_2_O, where spicule formation was only observed for RBCs interacting with ibuprofen molecules. In particular, higher ibuprofen concentrations (equivalent to 1200 mg and 2400 mg) caused RBC morphological changes that resulted in sphero-echinocytes that did not recover to normocytes, revealing a critical dose-dependent effect of ibuprofen and a potential implication for side effects concerning RBC health and function from overdosage (41).

Our results are consistent with previous findings based on SEM investigations, indicating progressive echinocytosis with increased ibuprofen concentrations (17). The morphological transition from a doughnut-like shape to an echinocyte morphology is suggested to originate from the interaction of the negatively charged ibuprofen particles with the RBC outer membrane bilayer, in accordance with the bilayer-couple hypothesis (Fig. 7*G-H*) (5, 42). An increase in the area between the inner and the outer monolayers of the RBC membrane, initiated by the binding of ibuprofen molecules, triggers echinocytosis (5). Higher concentrations of echinocytogenic compounds may result in a sphero-echinocyte RBC morphology, with a more distinct spherical shape and less pronounced spicules (43), which is consistent with our results. Higher sphericity and qualitatively less sharp specular structures were observed with higher ibuprofen concentrations (1.5-3 mM). Our findings highlight the dynamic formation and movement of single spicules on the membrane of ibuprofen-treated RBCs in real-time and provide evidence for a dose-dependent reversibility of RBC morphological alterations. The shape of the RBC is dependent on the interplay between the two main membrane components, which are the lipid bilayer and the spectrin cytoskeleton (44, 45). Thus, when the membrane asymmetry between the inner and the outer layers increases in favor of the outer layers, spicule formation is triggered as a natural response to the expansion of the outer leaflet coupled with the resistance of the cytoskeleton to the morphological distortion (44). The theoretical elastic membrane energy model and available experimental data support the preferential initial spicule formation on the RBC contour due to the highest curvature of the cytoskeleton (44). Driven by the continuous expansion of the outer monolayer, specular structures tend to move from the rim of the cell towards regions with a lower curvature, including the central area where the distinctive dimple is lost following the progression of echinocytosis, and finally distribute uniformly around the cell membrane (45). RBCs treated with higher ibuprofen concentrations showed that specular structures are more likely to steadily stay in place towards the later stages of echinocytosis, when a sphero-echinocyte morphology prevails. Before this occurs, the dynamic movement of spicules associated with increased membrane tension can induce spontaneous spicule splitting (45, 46). Here, a singular specular structure separates into two smaller daughter spicules as observed between 7:13 min and 7:36 min time points in Fig. 5*K-M*. Spicules are also seen dissolving (Fig. 5*N-P*) as the RBC shape returns to its discocyte morphology and the asymmetry between the two membrane leaflets is restored. Therefore, spicule motion tracking can provide real-time information on RBC nano-mechanics and it can act as a potential indicator for membrane bilayer defects (45). We suggest that in low ibuprofen concentration conditions, the rate of ibuprofen molecules interacting with the RBC membrane bilayer decreases over time, resulting in the transition back to a normocyte. With high ibuprofen concentrations, the constant interaction of ibuprofen molecules causes high RBC membrane asymmetry and the consequent inability of the sphero-echinocytes to recover their discocyte shape. Vesiculation and cell lysis are thought to occur at the final stages following spheroechinocytosis (44).

Alterations of the normal discocyte morphology of RBCs have a direct effect on RBC deformability, which determines not only the rheological properties but also the health and life span of single RBCs (6, 47). Echinocytosis presents a rheological disadvantage characterized by higher viscosity as well as decreased deformability, mainly driven by the increase in sphericity, with a direct impact on blood flow in large vessels and the ability of RBCs to squeeze through narrow capillaries, respectively (47, 48). The increased rigidity of echinocytes may also drive RBC aggregation, potentially contributing to a higher risk of occlusions of blood vessels and an impairment in the transport of oxygen (3, 44). The RBC shape changes reported in the present study and the associated alterations to the RBC morphological parameters, including a reduced surface area to volume ratio and an increased sphericity, are in agreement with a detrimental effect of ibuprofen on RBC rheological properties and overall health. Importantly, in our study, the inability of RBCs to recover their doughnut-like morphology was solely observed with high ibuprofen concentrations (1.5-3 mM), which correspond to 1200 mg and 2400 mg doses that should never be taken all at once. The most commonly used ibuprofen doses of 200 mg and 400 mg, corresponding to low ibuprofen concentration ranges used in the present study, showed a temporary echinocytosis progression. The widespread availability of ibuprofen as an OTC drug even at 1200 mg dosage increases the risk for overdosage and thus emphasizes the relevance of the observations reported in the present study in terms of drug safety. The potential risk from the continuous cumulative intake of standard ibuprofen doses over long periods of time, for instance for the treatment of rheumatoid arthritis, could not be assessed within the scope of this study.

In summary, the findings from our work highlight that the rheological properties of RBCs should be taken into account when formulating the safety levels for dose-dependent OTC and prescribed drugs intake, particularly NSAIDs. We anticipate that our ML-based label-free imaging approach operable with high spatial and temporal resolution even in resource-limited settings could be extended for detection of pathologies that can adversely affect RBC morphology, such as in neurocognitive disorders and transmissible diseases such as malaria (37, 49, 50).

## Materials and Methods

### Preparation of Blood Samples

Whole blood was freshly obtained with the consent of a healthy donor from a finger prick with safety lancets (VWR). Sickle cell trait (SCT) (ZenBio, SER-PRBC-AS) and sickle cell anemia (SCA) (BioIVT, HMRBC-SCKD) human red blood cell samples were commercially obtained from a single donor, respectively (*SI Appendix*, Table S1). For all blood samples, 10 μL of fresh blood was diluted in 10 mL PBS buffer (VWR) as a stock blood solution and 250 μL of the stock solution was transferred in a 35-mm Ibidi ibiTreat μ-Dish (Ibidi GmbH, Germany) for DHTM imaging. For AFM measurements, blood smears were prepared using 10 μL of fresh blood on SuperFrost glass slides (VWR) and were air-dried for 10 minutes.

### Preparation of Urea, H_2_O_2_ and ibuprofen solutions

Urea and H_2_O_2_ solutions were prepared by dissolving powder urea (~0.48 g/mL, Merk Millipore) and 30% H_2_O_2_ (Merk Millipore) in ddH_2_O, respectively. 2M, 4M, 6M and 8M stock solutions were prepared for both urea (*SI Appendix*, Fig. S6) and H_2_O_2_ (*SI Appendix*, Fig. S7). For each experimental condition, 50 μL of stock solution were added to 250 μL of stock blood solution in the petri dish after ~40s from the start of the live holo-tomographic video acquisition. The same volume of ddH_2_O alone was also tested as control (*SI Appendix*, Fig. S8). Ibuprofen powder was obtained by crashing a 400 mg ibuprofen tablet and stock solutions were prepared by dissolving ibuprofen (2 mg/mL) in 2 mL of PBS (VWR). Four concentrations of ibuprofen solution were prepared (0.25 mM, 0.5 mM, 1.5 mM and 3 mM). Based on the healthy donor weight of 60 kg and estimated blood volume of 65 mL/kg, the ibuprofen stock solutions corresponded to ibuprofen dosages of 200 mg, 400 mg, 1200 mg and 2400 mg and ibuprofen plasma concentrations of 51 μg/mL, 103 μg/mL, 308 μg/mL and 615 μg/mL (15). 50 μL of each ibuprofen stock solution was added to the blood cells as described above. For AFM measurements, 250 μL of healthy blood was incubated with 50 μL of each ibuprofen solution for 1.5 hours at 37°C, after which 10 μL was deposited on a glass slide and air-dried for 10 minutes. Additionally, 10 μL of ibuprofen solution (9.7 mM) was deposited on a gold thin film and air-dried overnight before imaging.

### Label-Free Digital Holo-Tomographic Microscopy

Label-free holo-tomographic imaging was performed using a 3D Cell Explorer microscope (Nanolive SA, Switzerland). During imaging, a top-stage incubator (Okolab srl, Italy) was used in order to control temperature (25°C), humidity and CO_2_. 4D RI tomograms were obtained at the highest temporal resolution of one frame every two seconds. Prior to each measurement, the petri dish containing the stock blood solution was placed inside the chamber of the top-stage incubator and the cells were allowed to sediment to the bottom of the petri dish for 10 minutes before imaging. For DHTM imaging of ibuprofen-treated blood smears, 25 μL of silicone oil (5 cSt, Merk Millipore) was added on the smear and a coverslip was placed on top and sealed with nail varnish (*SI Appendix*, Fig. S10).

### Atomic Force Microscopy

AFM measurements were performed on air-dried blood smear samples using the NaniteAFM with scan head 110 μm (Nanosurf AG, Switzerland). The glass slide was mounted onto the sample stage using the Nanite sample holder and the integral topview camera was used to locate a region of interest and to position it under the cantilever. The sideview camera was then used to perform an initial approach of the cantilever to the sample before the AFM final automatic approach. A Dyn190AI-10 AFM cantilever (Nanosurf AG, Switzerland) with self-alignment grooves, aluminium reflection coating, force constant 48 n/m and resonance frequency 190 kHz was used in phase contrast mode. Large-area 80 μm x 80 μm AFM images were obtained in order to identify non-overlapping RBCs, subsequently followed by single cell ~ 13 μm x 13 μm high-resolution AFM images. All AFM measurements were conducted at a scan rate between 0.5 Hz and 1.3 Hz. For imaging ibuprofen particles AFM measurements were conducted using multimode AFM (Bruker) using Scout 70 HAR Si tips (70 KHz, 2N/m) on ibuprofen particles deposited on gold thin films on mica substrate (Phasis Inc).

### Image Processing and Analysis

3D and 4D stacks obtained via DHTM were exported as TIFF files and imported into Imaris 9.8 (Bitplane AG, Switzerland). First, stacks were 3D cropped in the z-axis in order to include only slices that contained cells. Next, a 3×3×3 median filter was applied as a noise removal filter. Finally, a surface was fitted with background subtraction and automatic thresholding in order to achieve single cell segmentation. Additional filters were applied to the segmented image in order to filter out overlapping cells that could not be separated as well as partial cells touching the XY image borders. The morphologically-relevant features were quantitatively measured at the single cell level with Imaris, including the cell diameter, surface area, volume, thickness, sphericity and mean RI (*SI Appendix*, Table S3). The Hb concentration was calculated from the mean RI value of each single RBC, obtained from the 3D RI tomograms, using the following formula (51):

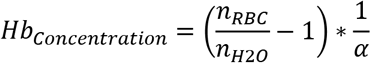

where *n_RBC_* is the mean RI value of the RBC, *n*_*H*2*O*_ is the RI of water (1.333) and α is the wavelength-dependent RI increment for RBCs, which was set to 0.001983 for λ = 520 nm (52). The Hb content was calculated for each single RBC by multiplying the *V* by the *Hb_Concentration_* (28). For 4D tomograms, the fitted surface was tracked during the entire duration of the time-lapse and the morphological features extracted for each individual frame. In order to benchmark the measurements for the morphological parameters with DHTM, we used micro-particles based on silicon dioxide (Merk Millipore) with diameters of 2 μm and 5 μm. The micro-particles were diluted in PBS, added to a glass slide and a coverslip was placed on top and sealed with nail polish in order to prevent drying. The morphological parameters were quantified with Imaris as described above and compared to the nominal values provided by the manufacturer (*SI Appendix*, Fig. S2). For the ML-based classification, Imaris ML feature based on Random Forest classification was used. A train-test data split of 33%-67% was applied for each experimental condition. During the training phase, single RBCs were manually annotated based on their morphology as normocyte, stomatocyte, echinocyte, acanthocyte, spherocyte, ovalocyte, helmet cell, sickle cell and teardrop cell (*SI Appendix*, Table S2). Next, the classifier predicted the morphology of the remaining cells based on the training data. All predictions were manually checked for accuracy in view of the low prevalence of some RBC types. For 4D tomograms, the ML-based classification was applied to each individual frame. For AFM image processing, the raw AFM data were analyzed using open source software Gwyddion 2.60. 2D levelling and scan line correction were applied before extraction of the height profile and surface roughness (RMS roughness, Sq) values. For the analysis of surface roughness distribution between healthy and ibuprofen-treated RBCs, a total of 250 single RBCs were analyzed for each sample. To calculate the size distribution of ibuprofen particles, a total of ~500 were analyzed.

### Molecular Dynamics Simulations. Modelling

The details of modelling RBC membrane bilayer with CHARMM-GUI (53, 54) based on the *in silico* lipid composition of the model erythrocyte membrane in ref. (55) is provided in Section S1.1 under Supplementary Information. Details of preparation of the five ibuprofen-lipid systems and molecular dynamics simulations with Gromacs 2018.4 (56) package using Charmm36m (57) force field to represent lipids and CHARMM General force field (58, 59) (CGenFF) to represent ibuprofen is provided in Section S1.2. Analyses of ibu–lipid and ibu–ibuprofen interaction energies, lipid headgroup mean square displacements (MSD) and diffusion coefficients (D), lipid hydrocarbon tail deuterium order parameters (S_CD_), and lipid heavy atom number density maps were performed by using *Gromacs tools*. The computed interaction energies plotted are normalized per ibuprofen molecule. The models were visualized using VMD (60).

## Acknowledgements

T.B. and P.N.N. thank Prof. Alex Dommann for helpful discussions and strategic support. This work used the computational resources provided by the University of Bern, Switzerland. T.B. thanks Guillaume Witz at the microscopy imaging center at University of Bern for technical assistance and Lars Lüder for graphical support. D.T. acknowledges support from Science Foundation Ireland (SFI) under award number 12/RC/2275_P2 (SSPC) and supercomputing resources at the SFI/Higher Education Authority Irish Center for High-End Computing (ICHEC).

