## Supplementary Information for "Label-free digital holotomography reveals ibuprofen-induced morphological changes to red blood cells"

### Section S1. Molecular dynamics model details

**S1.1 RBC membrane lipid bilayer model.** We modelled the RBC membrane lipid bilayer based on the *in silico* lipid composition of the model erythrocyte membrane in ref. (1), which have been previously experimentally validated. The membrane model was built using CHARMM-GUI (2, 3) web-interface and is composed of 20% and 20% cholesterol (CHL), 11% and 25% Palmitoyl Oleoyl PhosphoCholine (POPC), 38% and 10% Palmitoyl Oleoyl PhosphoEthanolamine (POPE), 22% and 0% Palmitoyl Oleoyl PhosphoSerine (POPS), and 9% and 35% Stearoyl SphingoMyelin (SSM) in the inner and outer leaflets, respectively (see Table S4 and Fig. S11*A*), with 200 lipid molecules in each leaflet. The surface area of the lipid bilayer was 10.5 X 10.5 $nm^2$ (Fig. S11*B*), large enough to study adsorption of ibuprofen aggregates on membrane surface.

**S1.2 Preparation of the ibuprofen-lipid systems and molecular dynamics simulations.** Molecular Dynamics (MD) simulations were performed using Gromacs 2018.4 (4) software. The ibuprofen molecules and RBC lipid bilayer were represented by CHARMM General force field (5, 6) (CGenFF) and CHARMM36m (7) force field parameters, respectively. *Five* different ibuprofen aggregates were studied on top of RBC membrane bilayer – *(1)* single molecule of ibuprofen (Fig. S11*C*), preformed aggregates of *(2)* 80 ibuprofen molecules, representing very low concentration, and *(3)* 100 ibuprofen molecules representing low concentration, but higher than 80-molecule aggregate, densely packed box of ibuprofen containing 1903 molecules under *(4)* isothermal-isobaric ensemble (NPT) conditions at constant pressure, where the volume of the system is adjusted during simulation representing high concentration, and *(5)* canonical ensemble (NVT) at constant volume to model very high concentration of ibuprofen aggregates. System *(1)* will henceforth be regarded as "single ibu", system *(2)* as "low ibu conc. I", system *(3)* as "low ibu conc. II", system *(4)* as "high ibu conc. I", and system *(5)* as "high ibu conc. II". The preformed aggregates of ibuprofen (80 and 100 molecules) were modelled by running a MD simulation of randomly dispersed molecules of ibuprofen in water with counterions ($Na^+$ and $Cl^-$) (see Fig. S11*D*). The ibuprofen aggregates formed instantly within 1 ns dynamics (Fig. S11*D*) due to strong hydrophobic intermolecular forces. The starting configuration of all five systems are shown in Figs. S12*A*–*E*. All ibuprofen–membrane complexes were solvated by filling the area above and below the membrane with water molecules represented by the modified TIP3P water model (8), creating a >20-Å thick water layer above the ibuprofen and below the membrane to mimic bulk solvation in the z-plane. Each simulation cell was neutralized by adding the appropriate number of counterions. After 5000 steps of energy minimization, each system was equilibrated over six consecutive steps (100 ps each), with the values of the force constants of position and dihedral restraints of lipids gradually decreased from 1000 to 0 (the unit for position and dihedral restraints are kJ/(mol.$nm^2$) and kJ/(mol.$rad^2$), respectively). During equilibration, the Berendsen thermostat and barostat were applied to maintain the temperature at 310 K and pressure at 1 atm. Semi-isotropic pressure coupling was applied to allow the lipid bilayer to fluctuate in the xy plane independent of the z-axis. For the production run, the Velocity rescaling thermostat and Parrinello-Rahman barostat were applied. Long-range electrostatic interactions were treated using the particle-mesh Ewald (PME) method. The time step used in our MD simulations is 2 fs, and the structures were saved every 100 ps during 0.1 µs of production dynamics.

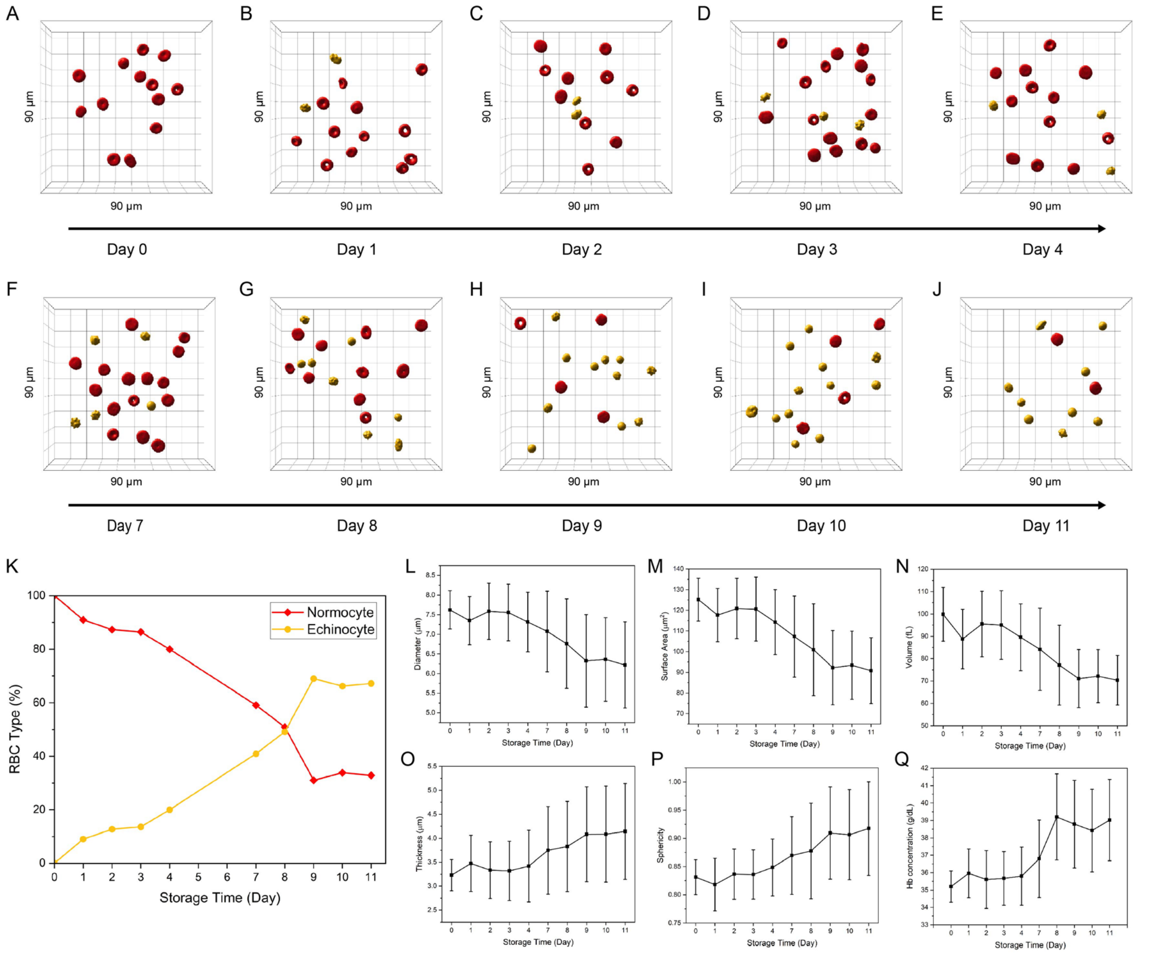


**Fig. S1.** Evolution of RBC quality over 11-days of storage time. 3D rendering of healthy RBCs diluted in PBS and stored at 4°C on (*A*) day 0, (*B*) day1, (*C*) day 2, (*D*) day 3, (*E*) day 4, (*F*) day 7, (*G*) day 8, (*H*) day 9, (*I*) day 10, (*J*) day 11. (*K*) Percentage of normocytes and echinocytes in the same blood solution over 11 days. (*L*)-(*Q*) Quantification of RBC morphological parameters over storage time: diameter, surface area, volume, thickness, sphericity and Hb concentration. Error bars depict the standard deviation. Field of view 90x90x30 µm.

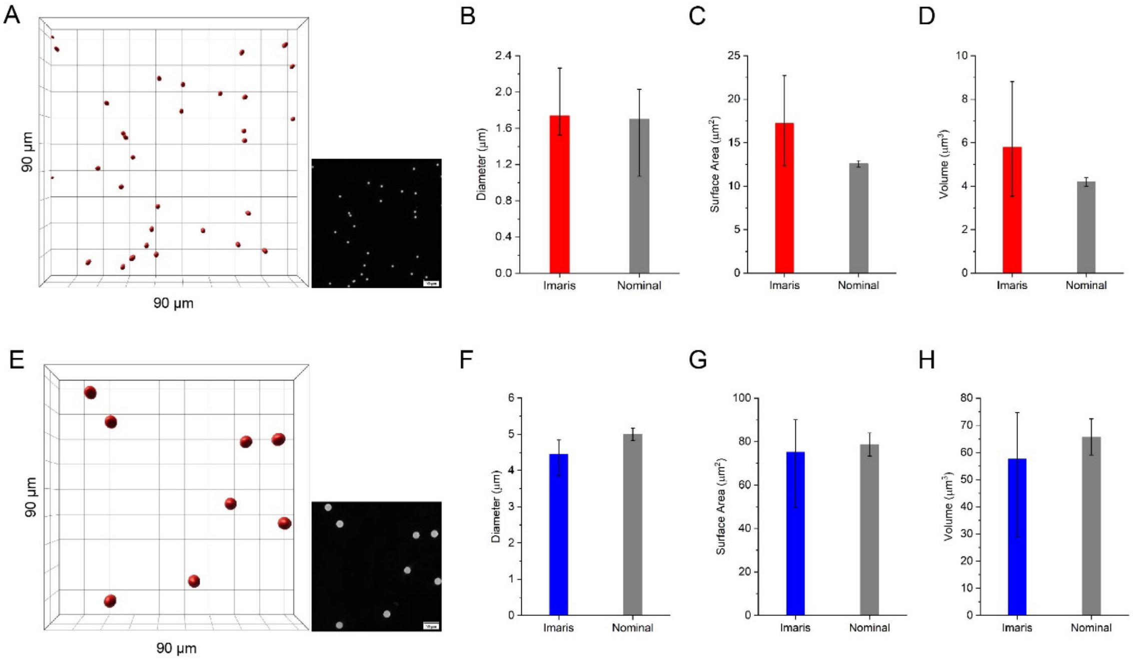


**Fig. S2.** Benchmark for the quantification of the morphological parameters using micro-particles based on silicon dioxide (Merk Millipore). (*A*) 3D rendering of 2 µm beads with corresponding RI tomogram in the inset, obtained with DHTM. Comparison of the quantification of (*B*) the diameter, (*C*) the surface area and (*D*) the volume, between the Imaris-based image analysis and the nominal values provided by the manufacturer. (*E*) 3D rendering of 5 µm beads with corresponding RI tomogram in the inset, obtained with DHTM. (*F*) Comparison of the quantification of the diameter, (*G*) the surface area and (*H*) the volume, between the Imaris-based image analysis and the nominal values provided by the manufacturer. Error bars depict the minimum and maximum values. Field of view 90x90x30 µm.

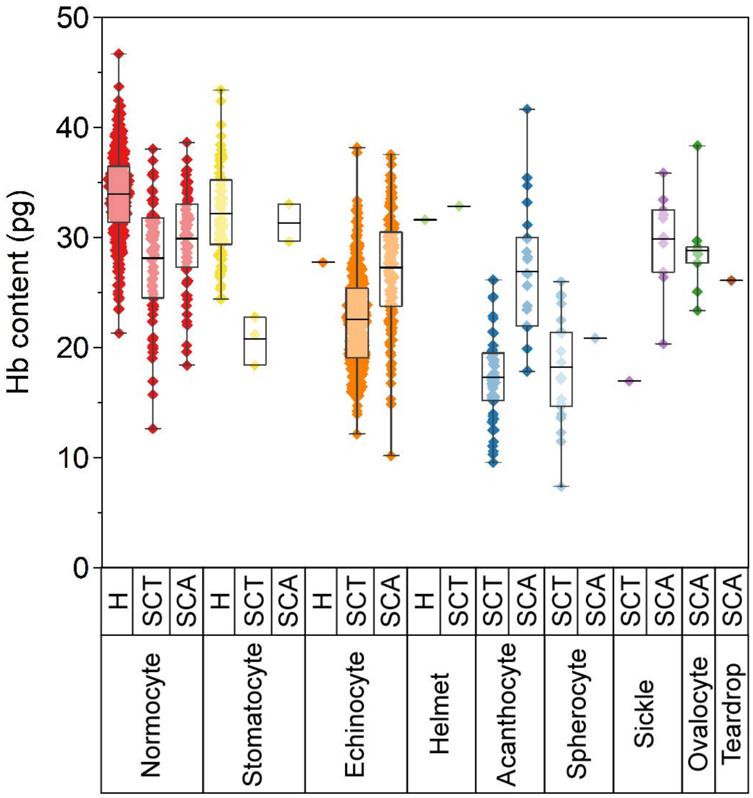


**Fig. S3.** Quantification of Hb content in healthy, SCT and SCA RBC populations based on 3D tomograms. Single cell level comparison between ML-based classified RBC types in healthy, SCT and SCA samples. Bars indicate mean values plus minimum and maximum values of all counted cells in each group.

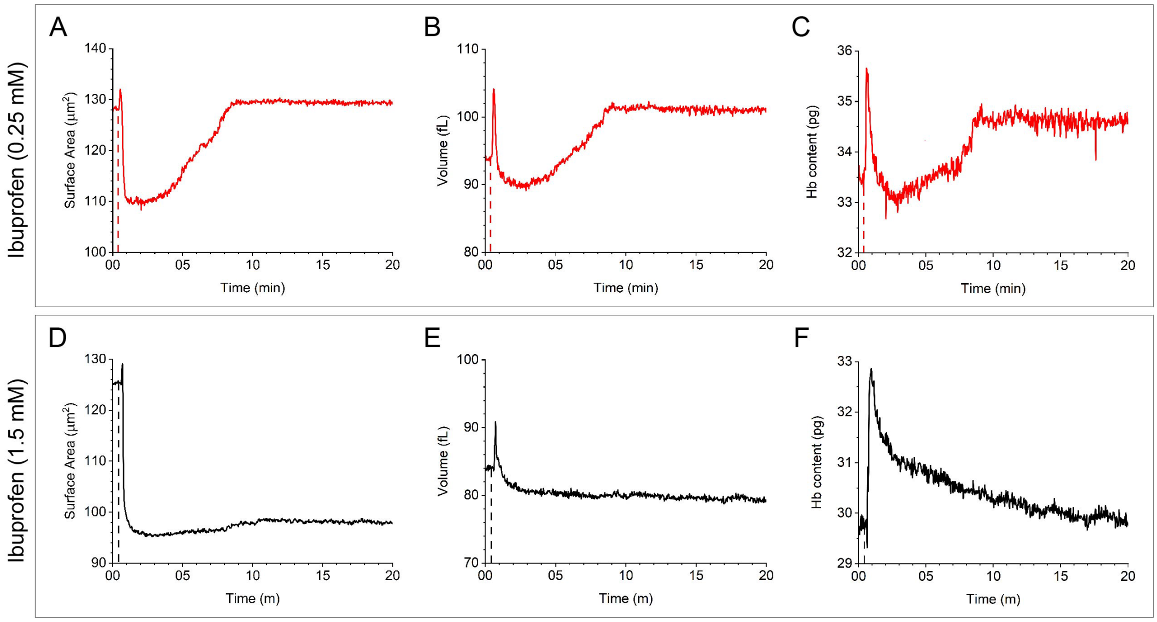


**Fig. S4.** Quantification of RBC morphological changes in surface area, volume and Hb content upon exposure to low and high concentrations of ibuprofen during a 20-minute time-lapse. (*A*-*C*) Time-dependent changes to surface area, volume and Hb content of a single RBC treated with 0.25 mM ibuprofen. (*D*-*F*) Time-dependent changes to surface area, volume and Hb content of a single RBC treated with 1.5 mM ibuprofen.

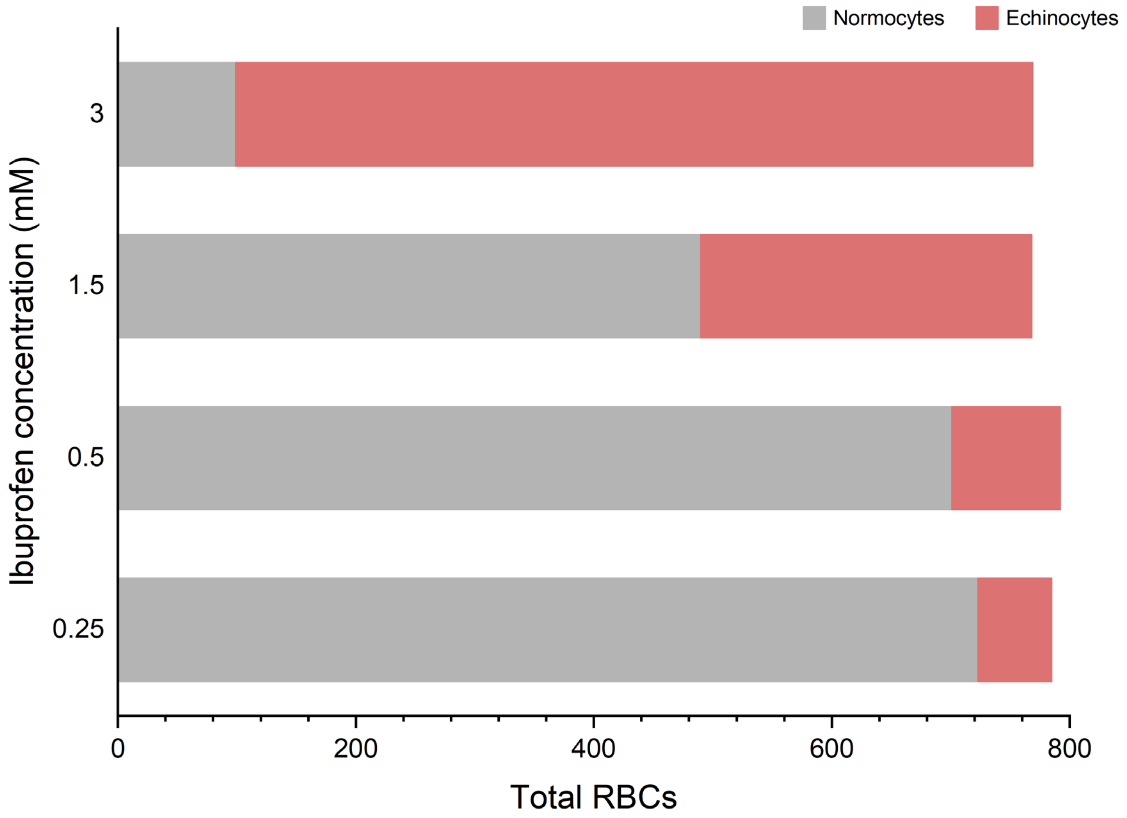


**Fig. S5.** Ratio of normocytes and echinocytes in blood incubated with 0.25 mM, 0.5 mM, 1.5 mM and 3 mM ibuprofen concentrations for 1.5 hours at 37°C.

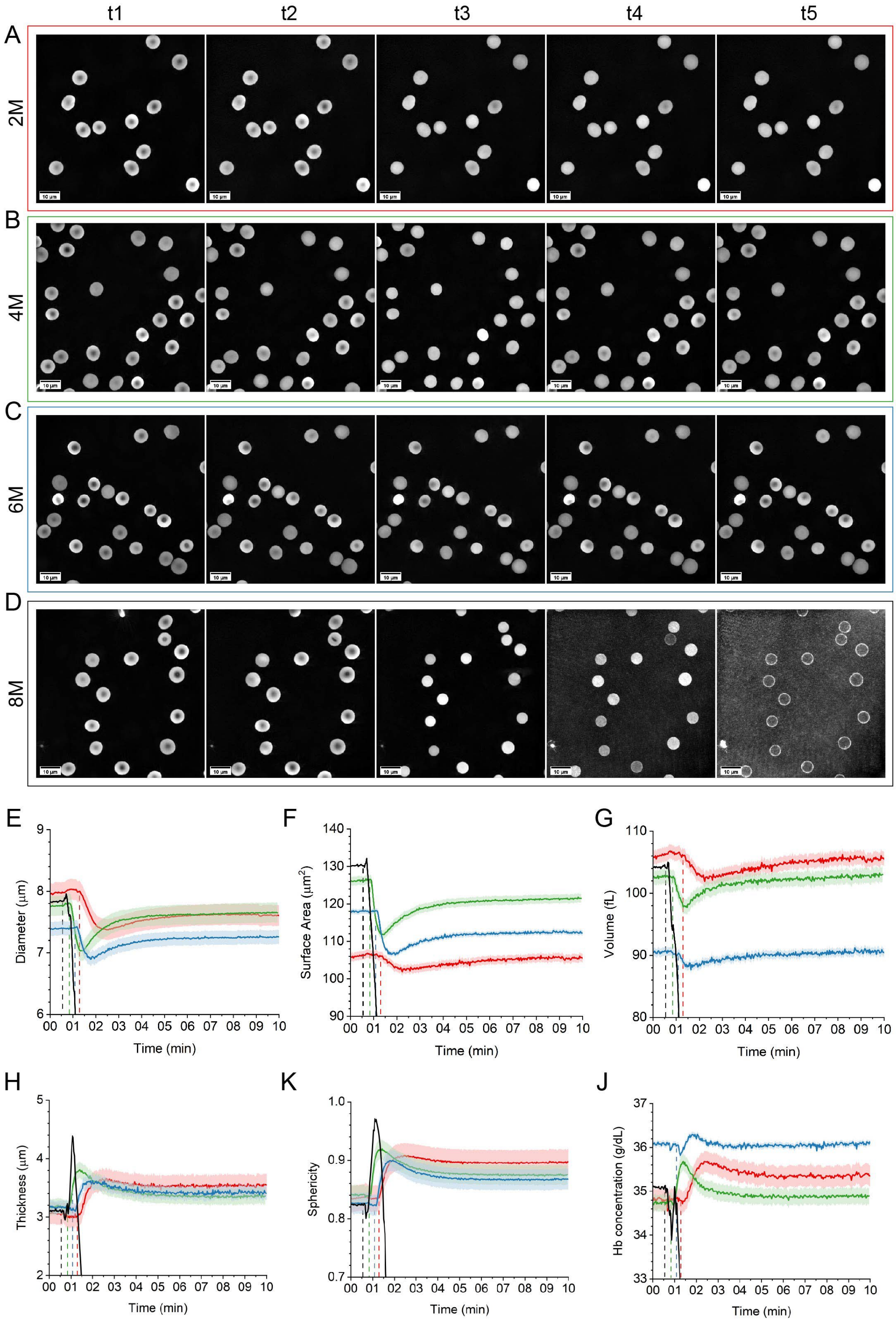
t1
t2
t3
t4
t5
A
2M
B
4M
C
6M
D
8M
E
Diameter (μm)
Time (min)
F
Surface Area (μm$^2$)
Time (min)
G
Volume (fL)
Time (min)
H
Thickness (μm)
Time (min)
K
Sphericity
Time (min)
J
Hb concentration (g/dL)
Time (min)

**Fig. S6.** 3D holo-tomographic imaging of RBCs treated with urea at varying concentrations, during a 10-minute time-lapse. (*A*) 3D RI tomograms of RBCs treated with 2M urea, resulting in mild spherocytosis (*t1*: 0 s; *t2*: 1:24 min, urea added; *t3*: 1:48 min; *t4*: 5:20 min; *t5*: 10 min). (*B*) 3D RI tomograms of RBCs treated with 4M urea, resulting in mild spherocytosis (*t1*: 0 s; *t2*: 1:02 min, urea added; *t3*: 1:26 min; *t4*: 4:58 min; *t5*: 10 min). (*C*) 3D RI tomograms of RBCs treated with 6M urea, resulting in mild spherocytosis (*t1*: 0 s; *t2*: 1:22 min, urea added; *t3*: 1:46 min; *t4*: 5:18 min; *t5*: 10 min). (*D*) 3D RI tomograms of RBCs treated with 8M urea, resulting in spherocytosis and cell lysis, with formation of ghost cells (*t1*: 0 s; *t2*: 48 s, urea added; *t3*: 1:08 min; *t4*: 2:20 min; *t5*: 10 min). (*E*-*J*) Quantification of time-dependent morphological parameters in urea-treated RBCs, (*E*) diameter, (*F*) surface area, (*G*) volume, (*H*) thickness, (*K*) sphericity and (*J*) Hb concentration (red = 2M; green = 4M; blue = 6M; black = 8M). Field of view 90x90x30 µm.

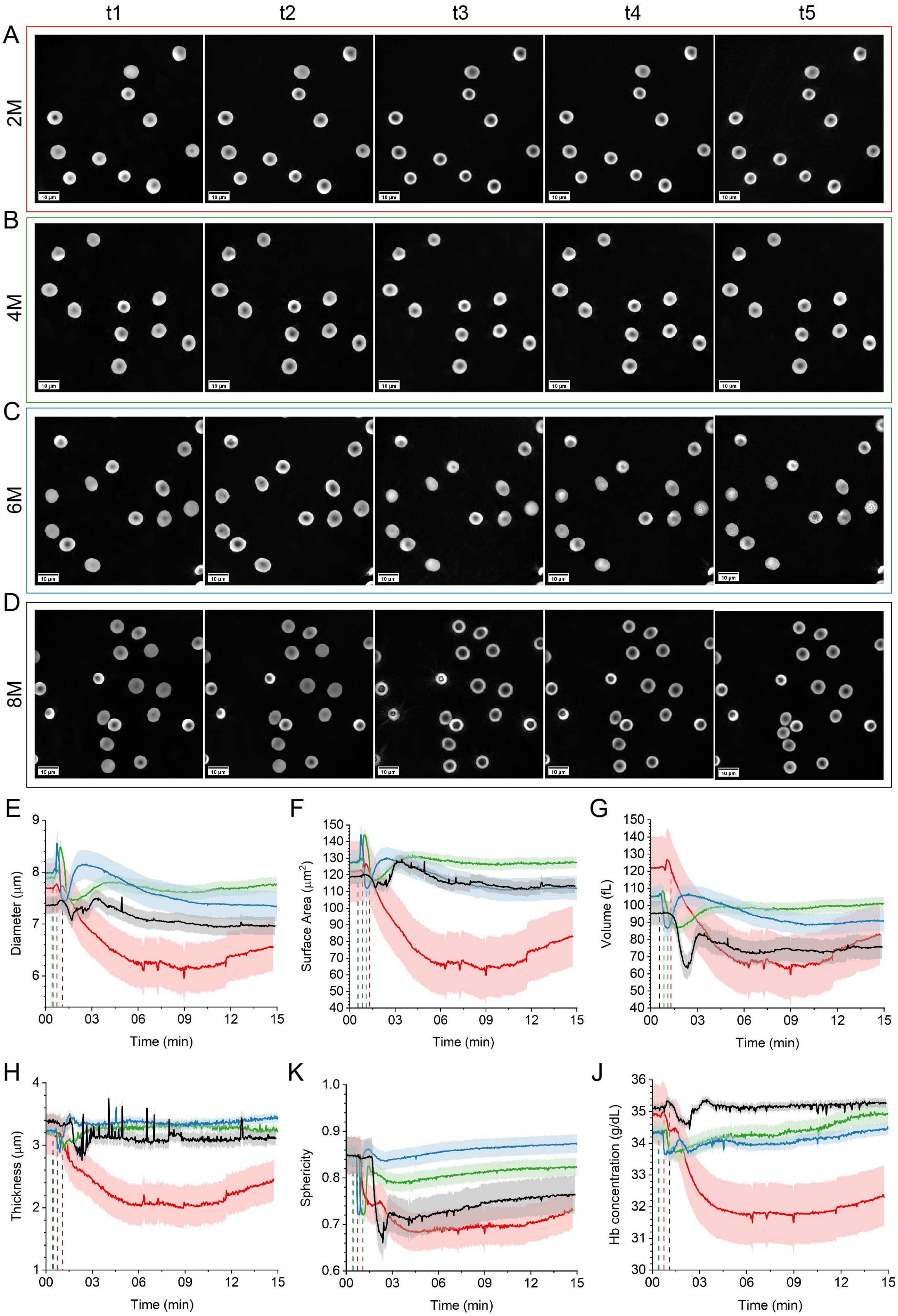

t1
t2
t3
t4
t5
A
2M
B
4M
C
6M
D
8M
10 µm
E
Diameter (µm)
Time (min)
F
Surface Area (µm²)
Time (min)
G
Volume (fL)
Time (min)
H
Thickness (µm)
Time (min)
K
Sphericity
Time (min)
J
Hb concentration (g/dL)
Time (min)

**Fig. S7.** 3D holo-tomographic imaging of RBCs treated with $H_2O_2$ at varying concentrations, during a 10-minute time-lapse. (*A*) 3D RI tomograms of RBCs treated with 2M $H_2O_2$, resulting in mild stomatocytosis (*t1*: 0 s; *t2*: 1:02 min, $H_2O_2$ added; *t3*: 1:48 min; *t4*: 7:44 min; *t5*: 15 min). (*B*) 3D RI tomograms of RBCs treated with 4M $H_2O_2$, resulting in mild stomatocytosis (*t1*: 0 s; *t2*: 54 s, $H_2O_2$ added; *t3*: 1:34 min; *t4*: 6:34 min; *t5*: 15 min). (*C*) 3D RI tomograms of RBCs treated with 6M $H_2O_2$, resulting in mild stomatocytosis (*t1*: 0 s; *t2*: 44 s, $H_2O_2$ added; *t3*: 1:28 min; *t4*: 6:24 min; *t5*: 15 min). (*D*) 3D RI tomograms of RBCs treated with 8M $H_2O_2$, resulting in more pronounced stomatocytosis (*t1*: 0 s; *t2*: 1:40 min, $H_2O_2$ added; *t3*: 2:20 min; *t4*: 7:20 min; *t5*: 15 min). (*E*-*J*) Quantification of time-dependent morphological parameters in urea-treated RBCs, (*E*) diameter, (*F*) surface area, (*G*) volume, (*H*) thickness, (*K*) sphericity and (*J*) Hb concentration (red = 2M; green = 4M; blue = 6M; black = 8M). Field of view 90x90x30 µm.

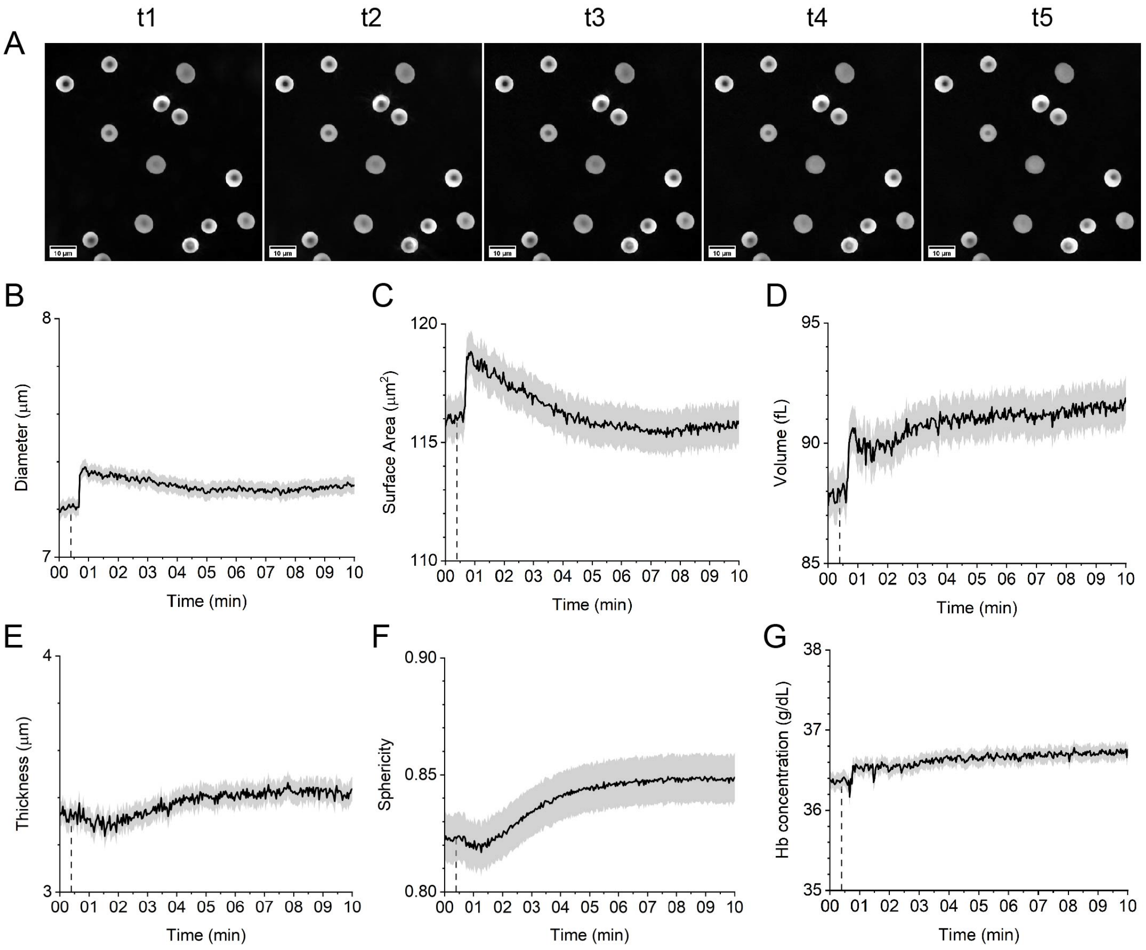


**Fig. S8.** 3D holo-tomographic imaging of RBCs treated with $ddH_2O$, during a 10-minute time-lapse. (*A*) 3D RI tomograms of RBCs treated with $ddH_2O_2$, resulting in no morphological alteration (*t1*: 0 s; *t2*: 44 s, $ddH_2O$ added; *t3*: 1:04 min; *t4*: 4:00 min; *t5*: 10 min). (*B*-*G*) Quantification of time-dependent morphological parameters in urea-treated RBCs, (*B*) diameter, (*C*) surface area, (*D*) volume, (*E*) thickness, (*F*) sphericity and (*G*) Hb concentration. Field of view 90x90x30 µm.

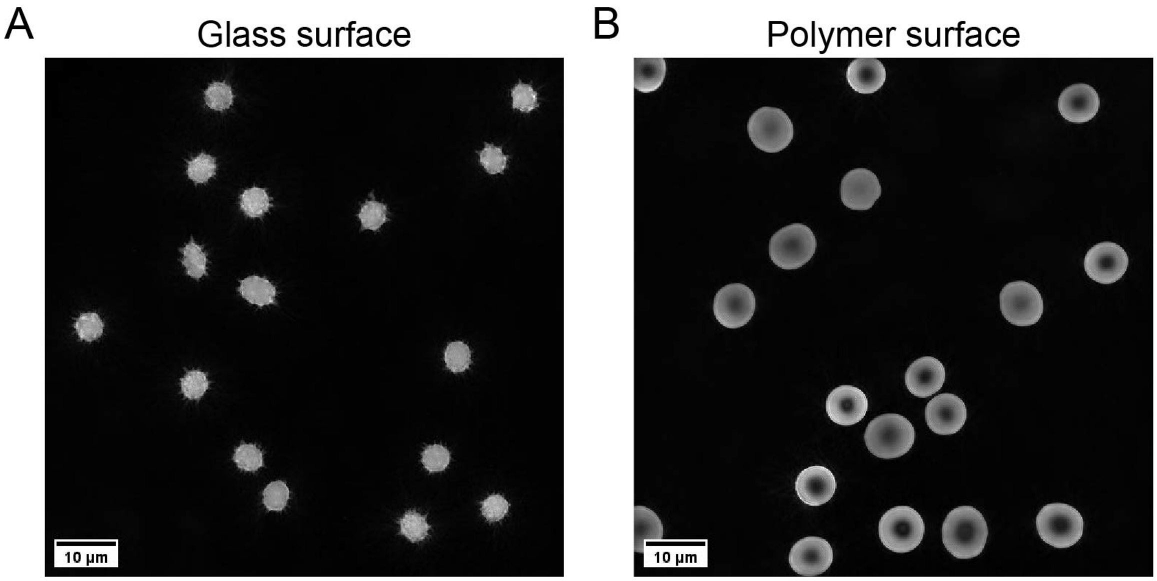


**Fig. S9.** Effect of petri dish surface used to image blood solutions with DHTM. (*A*) Echinocytosis resulting from RBCs contact with a glass surface. (*B*) Polymer-coated surface resulting in unaltered RBC morphology.

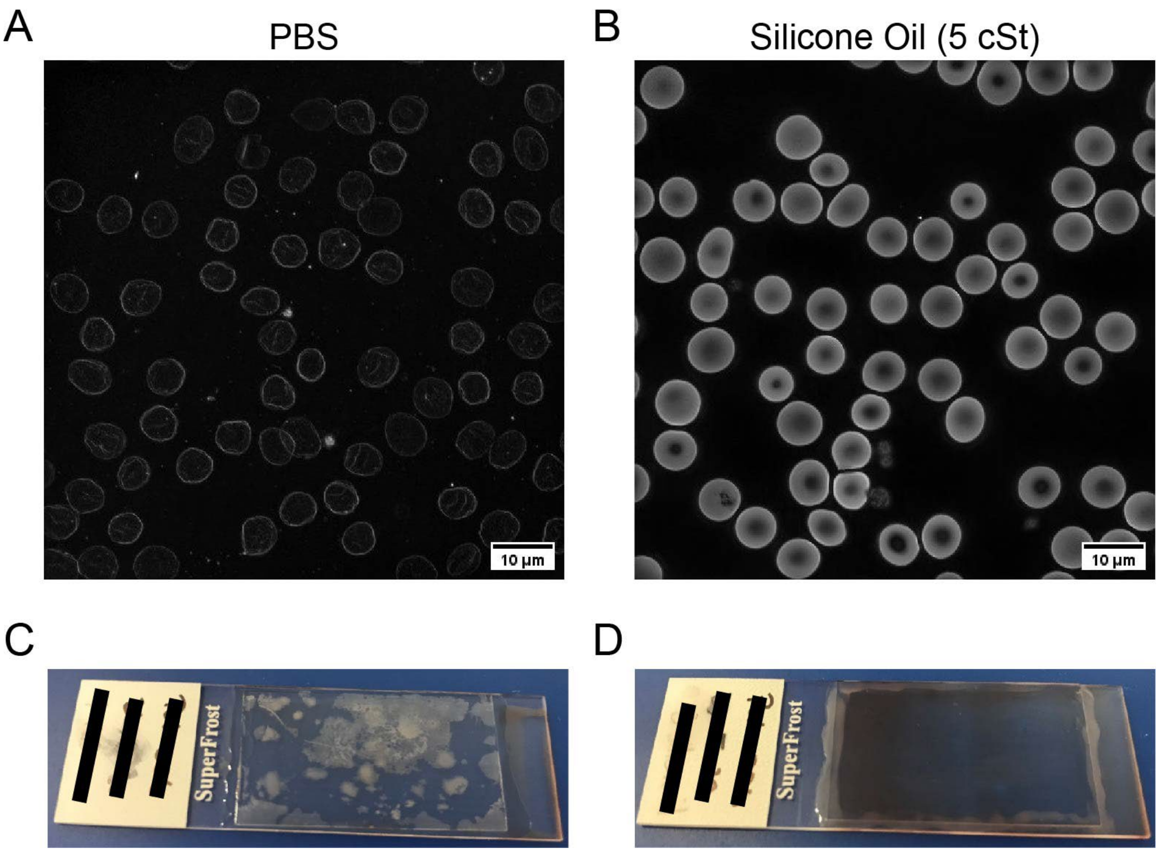


**Fig. S10.** Comparison between PBS and silicone oil (5 cSt) coating for optimal blood smear imaging with DHTM. (*A*) RI tomogram of healthy RBCs from a blood smear coated with PBS. (*B*) RI tomogram of healthy RBCs from a blood smear coated with silicone oil (5 cSt). (*C*) Example of storage artifacts in a blood smear coated with PBS and sealed with a coverslip and nail varnish after 4 days storage at room temperature. (*D*) Example of absence of storage artifacts in a blood smear coated with silicone oil (5 cSt) and sealed with a coverslip and nail varnish after 4 days storage at room temperature.

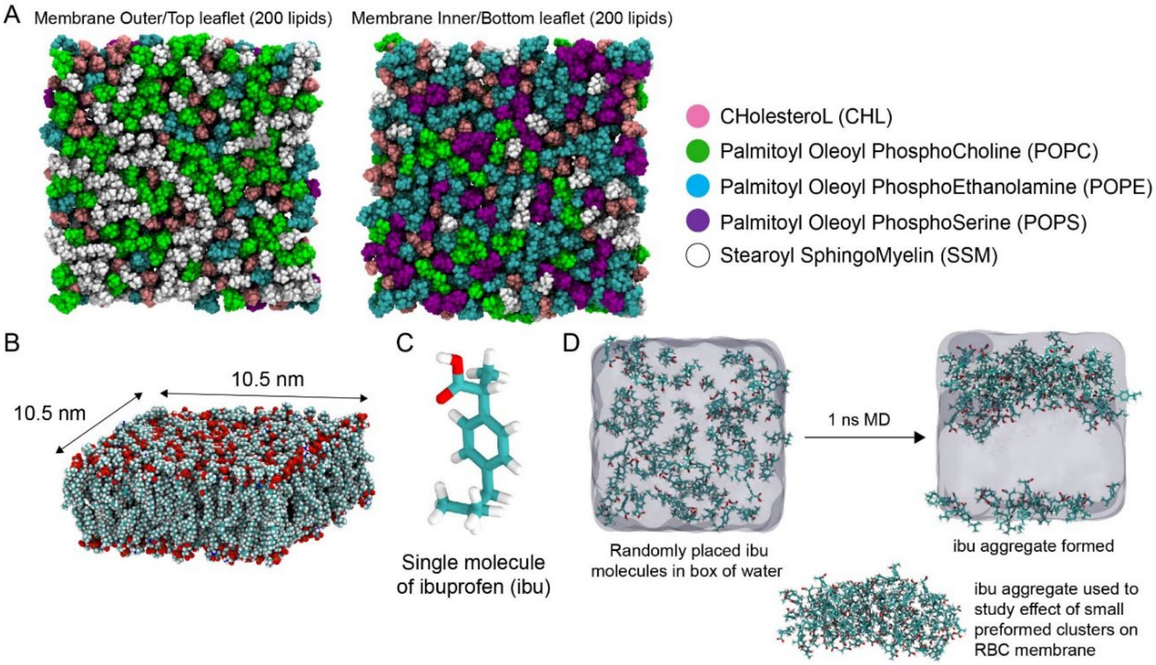


**Fig. S11.** *(A)* Model RBC lipid bilayer composition at the outer/top and inner/bottom leaflets used in this study also showing the *(B)* membrane dimensions in the xy-plane. *(C)* Stick representation of a single molecule of ibuprofen (ibu). *(D)* Preformed aggregates of ibuprofen molecules (at low concentration) obtained in this study from 1 ns dynamics in explicit water.

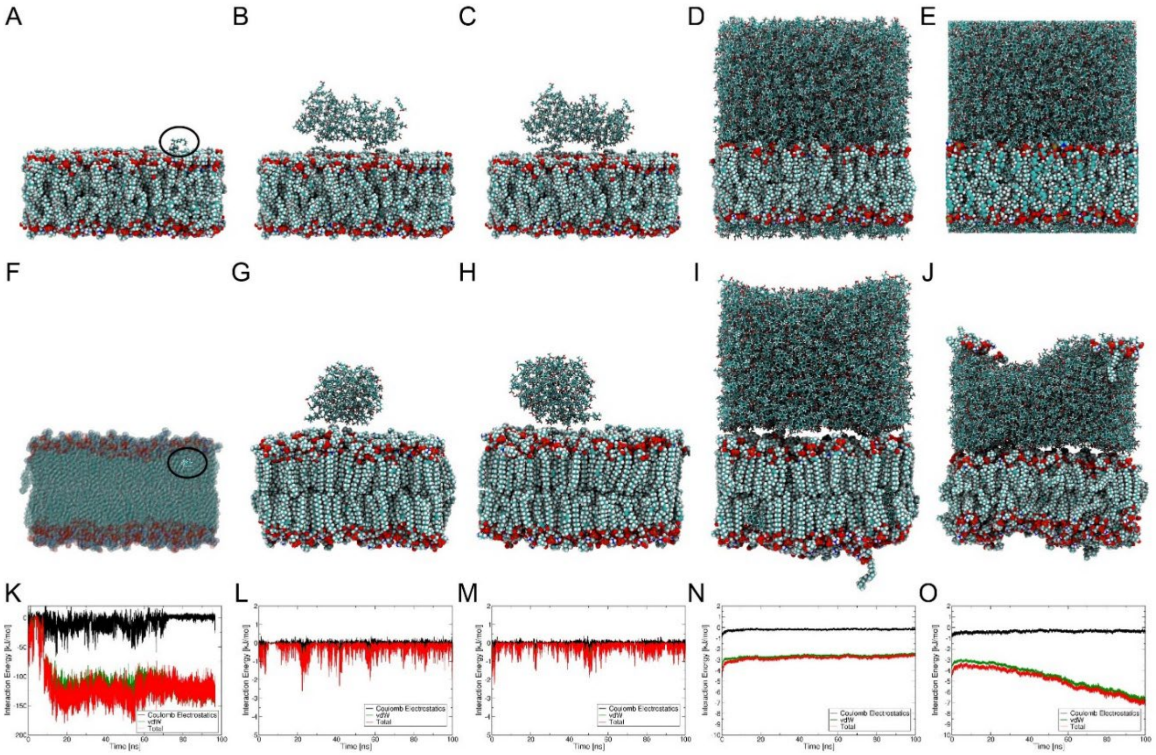


**Fig. S12.** *(A-E)* Initial and *(F-J)* final snapshots of *(A, F)* single ibuprofen molecule, aggregates of *(B, G)* 80 ibuprofen molecules, *(C, H)* 100 ibuprofen molecules, densely packed *(D, I)* 1903 ibuprofen molecules under NPT conditions, and *(E, J)* 1903 ibuprofen molecules under NVT conditions. *(K-O)* Ibuprofen–membrane interaction energies and their Coulombic electrostatic and van der Waals (vdW) components for all systems modelled in this study.

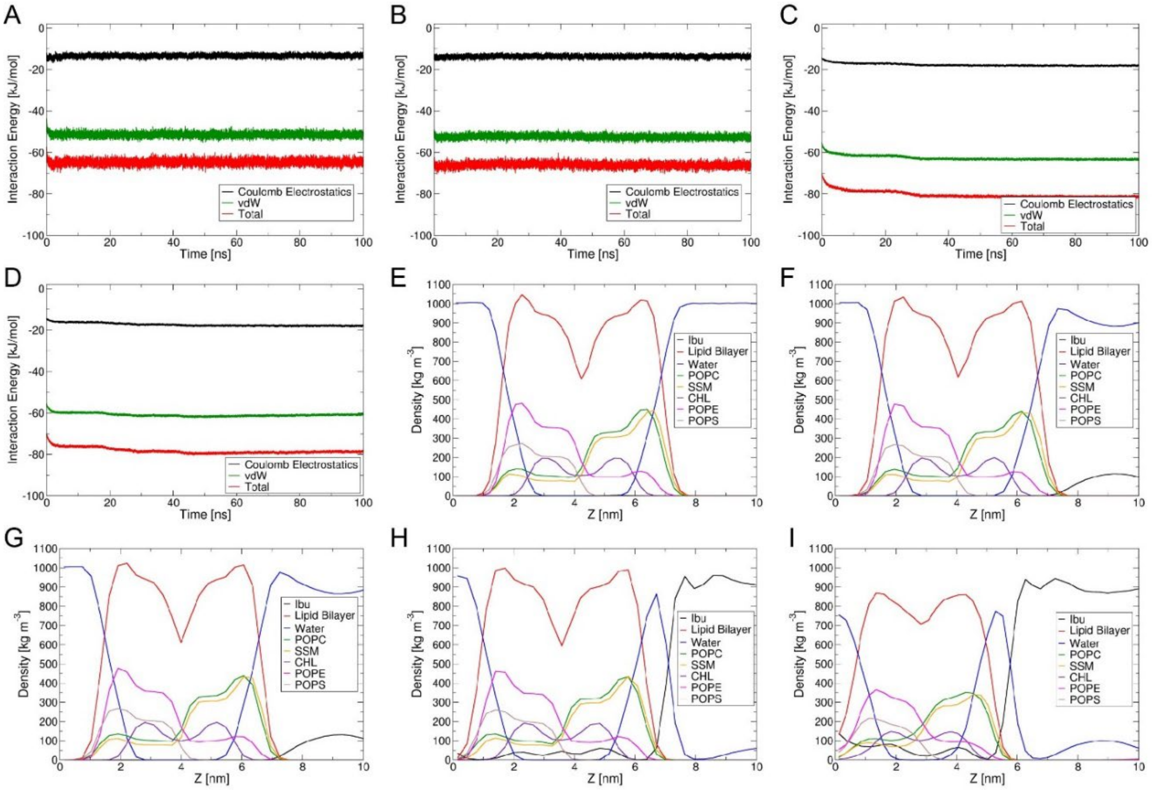


**Fig. S13.** Ibuprofen–ibuprofen interaction energies and their electrostatic and vdW components for *(A)* low conc. I, *(B)* low conc. II, *(C)* high conc. I, and *(D)* high conc. II (see Section S1.2 for these definitions). *(E-I)* Average density profiles of all species in the simulation box for all systems used in this study.

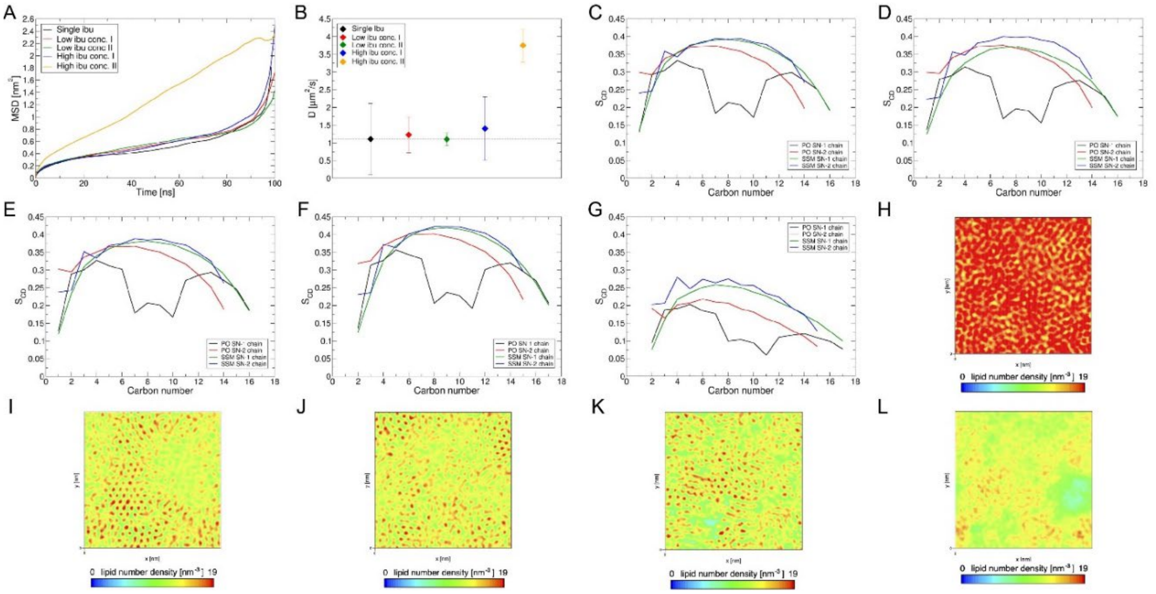

**Fig. S14.** Comparison of *(A)* Mean Square Displacement (MSD) and *(B)* diffusion coefficient, *D* for all systems modelled in this study. (C-G) Lipid hydrocarbon tail deuterium order parameter ($S_{CD}$) for RBC membrane lipids (Palmitoyl Oleoyl and Stearoyl) with two acyl chains, SN-1 and SN-2, and (H-L) lipid heavy atoms number densities in the *xy*-plane averaged over the *z*-axis to obtain a top view of lipid densities in the membrane for all systems.

**Table S1.** Summary of the demographics of the healthy blood and SCT and SCA RBC samples.

| | **Gender** | **Age** | **Ethnicity** |
|---|---|---|---|
| **Healthy** | F | 27 | Caucasian |
| **SCT** | M | 34 | Caucasian |
| **SCA** | F | 28 | African American |

**Table S2.** Overview of the total number of analyzed RBCs for the healthy, SCT and SCA samples and the percentages of each RBC type that was present in each sample.

| | **Total number of cells** | **RBC type** | | | | | | | | |
|---|---|---|---|---|---|---|---|---|---|---|
| | | Normocyte | Stomatocyte | Echinocyte | Helmet | Acanthocyte | Spherocyte | Sickle | Ovalocyte | Teardrop |
| **Healthy** | 351 | 272 (77.5%) | 77 (21.9%) | 1 (0.3%) | 1 (0.3%) | 0 (0%) | 0 (0%) | 0 (0%) | 0 (0%) | 0 (0%) |
| **SCT** | 459 | 71 (15.5%) | 0 (0%) | 306 (66.7%) | 1 (0.2%) | 55 (12%) | 25 (5.4%) | 1 (0.2%) | 0 (0%) | 0 (0%) |
| **SCA** | 230 | 62 (27%) | 2 (0.9%) | 122 (53%) | 0 (0%) | 23 (10%) | 1 (0.4%) | 10 (4.3%) | 9 (4%) | 1 (0.4%) |

**Table S3.** Description of the extracted morphological parameters from Imaris 9.7.

| Morphological Parameter | Description |
|---|---|
| Diameter | The length of the longest principal axis inside the object (*BoundingBoxOO Length C*) |
| Surface Area | The sum of the triangle surfaces |
| Volume | Quantification of how much a surface object occupies |
| S/V Ratio | Surface area divided by the volume |
| Thickness | The length of the shortest principal axis (*BoundingBoxOO Length A*) |
| Sphericity | The ratio of the surface area of a sphere to the surface area of the particle |
| Mean RI | Mean intensity of voxels enclosed within the surface |

**Table S4.** Composition of RBC membrane lipid bilayer molecular model. CHL = Cholesterol, POPC = Palmitoyl Oleoyl PhosphoCholine, POPE = Palmitoyl Oleoyl PhosphoEthanolamine, POPS = Palmitoyl Oleoyl PhosphoSerine, and SSM = Stearoyl SphingoMyelin.

| | CHL | POPC | POPE | POPS | SSM |
|---|---|---|---|---|---|
| **Inner leaflet** | 40 (20%) | 22 (11%) | 76 (38%) | 44 (22%) | 18 (9%) |
| **Outer leaflet** | 40 (20%) | 70 (35%) | 20 (10%) | 0 (0%) | 70 (35%) |

**Movie S1.** Effect of 0.25 mM ibuprofen on RBCs. RI tomograms were acquired at 2 sec intervals over a period of 20 min. 3D renderings of extracted frames from the video are provided in Fig. 3. Timecode is min:sec.

**Movie S2.** Effect of 0.5 mM ibuprofen on RBCs. RI tomograms were acquired at 2 sec intervals over a period of 20 min. 3D renderings of extracted frames from the video are provided in Fig. 3. Timecode is min:sec.

**Movie S3.** Effect of 1.5 mM ibuprofen on RBCs. RI tomograms were acquired at 2 sec intervals over a period of 20 min. 3D renderings of extracted frames from the video are provided in Fig. 3. Timecode is min:sec.

**Movie S4.** Effect of 3 mM ibuprofen on RBCs. RI tomograms were acquired at 2 sec intervals over a period of 20 min. 3D renderings of extracted frames from the video are provided in Fig. 3. Timecode is min:sec.

**Movie S5**. 3D segmented rendering of a single RBC exposed to low (0.25 mM) ibuprofen concentration and measured with DHTM, showing transient spicule formation, movement and dissolution across the RBC membrane. Timecode is min:sec.

**Movie S6**. 3D segmented rendering of a single RBC exposed to high (1.5 mM) ibuprofen concentration and measured with DHTM, showing irreversible spicule formation and movement across the RBC membrane. Timecode is min:sec.
